# Dynamic structural equation models for ecosystem-based fisheries management: case studies and a practical guide

**DOI:** 10.64898/2026.09.05.749621

**Authors:** Jennifer S. Bigman, James T. Thorson, Scott I. Large, S. Kalei Shotwell, Cole C. Monnahan, Silvana Gonzalez, Sarah Wise

## Abstract

Once central challenge in adopting ecosystem-based fisheries management (EBFM) is understanding and predicting the interconnected processes, feedbacks, and temporal dependencies that characterize ecosystem dynamics. Structural equation models and structural causal models (SEM and SCM) quantify interconnected relationships and evaluate causality from observational data but cannot readily accommodate the incomplete and irregularly sampled data typical of complex systems. Dynamic structural equation models (DSEM) estimate SEMs but accommodate lagged effects, missing data, and non-linear and non-stationary effects. However, there is little guidance regarding how DSEM can be used to support EBFM, particularly within governance and decision-making contexts. Here, we present three case studies that highlight distinct inferential goals for understanding ecosystem processes in the context of EBFM. First, we quantify ecosystem-level indicators to support monitoring of climate-driven ecosystem changes. Second, we examine whether hypothesized direct and indirect ecosystem linkages improve predictions of recruitment. Finally, we compare support for hypotheses of causal drivers of fish productivity. To provide practical advice, we synthesize lessons learned into a workflow to for others navigating DSEM. We suggest that the first step is identifying the inferential objective, which establishes the foundation for the remainder of the workflow, guiding the development of graphical models, model specification, and the selection of statistical and scientific validation steps (e.g., testing conditional independencies) needed to evaluate hypothesized causal relationships. By integrating observational data, expert knowledge, and mechanistic hypotheses within a unified framework, DSEM is a powerful tool for advancing EBFM, and more broadly, understanding the processes that govern complex ecosystems.

## Introduction

Fisheries management aims to understand the dynamics of, and sustainably manage, living marine resources. Currently, management practices in the United States and elsewhere focus on maximizing the harvest of a single species based on abundance, biology, and catch (Methot et al., 2014; Möllmann et al., 2014; Skern-Mauritzen et al., 2016). In the United States, the dynamics of individual stocks are modeled over time and translated into harvest policies and catch advice through collaboration with fishery councils and other groups that are composed of scientists, fishers, industry partners, tribal members, and other experts (Lynch et al., 2018)8/6/2026 11:02:22 AM. This single-species framework does not explicitly include the many factors that affect population dynamics including ecological interactions, environmental variability, socioeconomic factors that affect fisher behavior, incentives, and adaptive decision-making under conditions of uncertainty, among others (Howell et al., 2021; Marshall et al., 2019; Pikitch et al., 2004). Although such a management approach can attribute changes in abundance to processes such as recruitment or fishing mortality, it often provides limited insight into broader ecological and ecosystem drivers, complicating our ability to sustainably manage fisheries (Brewster et al., 2025; Holsman et al., 2020; Murawski, 1991).

Ecosystem-based fisheries management (EBFM) is a paradigm in which fisheries and other marine resources are managed holistically by incorporating many aspects of an ecosystem in the management process (Link, 2002; Pikitch et al., 2004). Such an approach is expected to ameliorate many of the challenges of more traditional, single-species fisheries management. For example, multispecies stock assessment models account for the effects of predator–prey relationships on individual stocks simultaneously and are one of the tools for EBFM (Holsman et al., 2020; Karp et al., 2023; Trijoulet et al., 2020). Work has shown that not accounting for predation mortality in stock assessments results in higher uncertainty in estimated stock biomass (Hollowed et al., 2020). Another example is the effect that environmental variability has on stock dynamics such as growth, mortality, and recruitment (Lee et al., 2012; Maunder & Watters, 2002; Skagen et al., 2013). Simulations have demonstrated that including variation in somatic growth— which varies with the environment and other ecosystem processes—led to increased precision and less bias when estimating management-related quantities such as stock spawning biomass (Correa et al., 2021). However, few operational stock assessments (i.e., those used for management advice) across the U.S. and elsewhere directly account for environmental and ecosystem considerations (Karp & Vieser, 2024; Marshall et al., 2019; Pepin et al., 2022; Skern-Mauritzen et al., 2016).

Although stock assessments form the basis of fisheries management, there are numerous pathways to incorporate ecosystem information into the management process that help us shift towards an ecosystem-based approach (Limpinsel et al., 2025; Shotwell et al., 2023; Townsend et al., 2019; Zador et al., 2017). For example, NOAA Fisheries produces annual Ecosystem Status Reports that assess the status and trends of different environmental/oceanographic, biological, and socioeconomic factors (Gove et al., 2022; Siddon, 2024). Such reports facilitate implementation of EBFM as the information can provide critical context and contribute to the setting of harvest recommendations (Dorn & Zador, 2020; Zador et al., 2017). Similarly, Ecosystem and Socioeconomic Profiles (ESPs) focus on individual stocks by summarizing and evaluating the importance of ecosystem and socioeconomic drivers of stock dynamics, thereby laying the groundwork for incorporating these factors directly into stock assessments (Shotwell et al., 2023). Ecosystem models, which range from qualitative (e.g., network models that provide a simple avenue to explore ecosystem tradeoffs) to complex (trophic models that incorporate mass-balance and predator-prey interactions), have also been used to support the integration of ecosystem information into the management process (Collie et al., 2016; Rodriguez-Perez et al., 2023; Townsend et al., 2019). Additionally, management systems can formally embed ecosystem information into management decisions. For example, ecosystem information within the context of the Ecosystem Status Report and the risk table framework (Dorn & Zador, 2020) helped explain the drivers of the Gulf of Alaska Pacific cod stock decline following a marine heatwave, while the management process fostered communication among ecosystem scientists, stock assessment scientists, managers, and industry stakeholders, facilitating the integration and acceptance of this information into catch-setting decisions (Barbeaux et al., 2020; Townsend et al., 2019; Zador & Yasumiishi, 2017).

Disentangling how complex ecosystem processes affect population dynamics—including recruitment, growth, and mortality—remains a major scientific barrier to implementing ecosystem approaches to management (Audzijonyte et al., 2025; Frid et al., 2006; Xu et al., 2025). Climate-driven changes in fish biology can mimic the effects of fishing mortality (Audzijonyte et al., 2014; Waples & Audzijonyte, 2016), and relationships between population dynamics and ecosystem variables can be non-stationary, meaning they vary across time and space (Wainwright, 2021; Ward et al., 2022). Such variability is challenging to understand and complicates efforts to implement more holistic management. For example, the Pacific sardine stock assessment model included a sea surface temperature effect on recruitment for several years before it became evident that the relationship no longer held (due to autocorrelation, see Jacobson & MacCall, 1995; McClatchie et al., 2010). This relationship was originally identified using a generalized additive model, which, like other regression approaches, primarily identifies statistical associations rather than causal or mechanistic relationships (Carlin & Moreno-Betancur, 2025; Grace & Irvine, 2020). Moreover, regression models can be sensitive to confounding bias—where an external variable affects both the response and predictor—and to collinearity among predictors, both of which can bias estimates of effect sizes (Arif & MacNeil, 2023; Byrnes & Dee, 2025; Grace & Irvine, 2020). Consequently, regression-based approaches may be poorly suited to identifying the mechanisms by which environmental variability influences fisheries dynamics (Arif & MacNeil, 2023; Champagnat et al., 2026; Thorson et al., 2024). However, the regression approach remains popular due to the lack of a scientific and statistical framework capable of simultaneously describing and predicting ecosystem dynamics, forecasting future states, and identifying causal drivers of population change needed for more complete ecosystem understanding.

Structural equation modeling (SEM) and structural causal modeling (SCM) are increasingly being used to understand and predict the dynamics of complex systems and evaluate mechanistic hypotheses using observational data (Byrnes & Dee, 2025; Correia et al., 2026; Grace et al., 2010; Grace & Irvine, 2020). SEMs are generally descriptive and focus on estimating the strength and direction of relationships among variables in the assumed graphical model represented as a directed acyclic graph (DAG; Arif & MacNeil, 2023). SCM is a broader framework for causal inference that uses DAGs to explicitly represent causal assumptions and evaluate whether observed data are consistent with hypothesized cause-and-effect mechanisms (Arif & MacNeil, 2023; Byrnes & Dee, 2025; Grace et al., 2010; Grace & Irvine, 2020). These approaches have become powerful tools for ecologists working with observational data in settings outside fisheries science and, specifically, EBFM. For example, SEM and SCM have been used to uncover environmental factors affecting earthworm diversity (Goury et al., 2025), understand whether species richness changes productivity (Dee et al., 2023), and identify the effect of environmental conditions and demography on coral reef recovery post-disturbance (Gouezo et al., 2019). However, many ecological systems are dynamic through time and relationships among variables often involve both simultaneous and lagged effects. Representing these temporal dependencies can be challenging using traditional SEM and SCM frameworks. Dynamic structural equation modeling (DSEM) is an approach for estimating SEMs that accommodates lagged effects, missing data, and non-linear and non-stationary effects, making it particularly useful for ecological time series (Fu et al., 2026; Thorson et al., 2024). For example, Champagnat et al. (2026) integrated DSEM into a stock assessment model to explain variation in walleye pollock (*Gadus chalcogrammus*) recruitment with observed environmental and ecosystem time series, improving the ability to explain past recruitment trends and project future dynamics. Despite its potential in uncovering how ecosystem dynamics may affect fisheries, there remain few examples of how DSEM can be used to facilitate EBFM.

To that end, we highlight how DSEM can be used in the context of EBFM to account for the complex, interrelated nature of ecosystem dynamics and how they may affect biological processes related to fisheries management. We focus on three case studies that each have a distinct scientific inferential goal, or the type of conclusion the analysis is intended to support (e.g., description, prediction, causal inference). Our first case study illustrates how DSEM can be used to estimate broad-scale, ecosystem-level trends by quantifying latent dynamics of benthic and pelagic taxa in the Chukchi Sea. Second, we examine whether hypothesized direct and indirect ecosystem linkages improve predictions of Gulf of Alaska arrowtooth flounder recruitment, a species responsible for intense predation on many other commercially important groundfishes in the region. The final case study uses DSEM to evaluate and compare support for three mechanistic hypotheses driving productivity of fishes in the Northwest Atlantic: whether juvenile abundance is driven by the condition and abundance of spawners, prey availability, or both. These case studies were chosen to demonstrate the breadth of possible applications of DSEM to EBFM, but also how the desired inferential goal impacts the DSEM workflow, including how to formulate causal hypotheses as DAGs, select a method to validate the models scientifically and statistically, and interpret the results. Finally, we synthesize our findings into a workflow diagram to guide others in integrating observational data, expert knowledge, and mechanistic hypotheses to better understand complex ecological systems using autocorrelated, patchy data.

Case Studies

Below we present the background, hypothesis and inferential goal, data, model(s), and results for each of our case studies separately. These analyses are intended to demonstrate different workflows that could be adopted and adapted for using DSEM for specific inferential goals and research questions. All data and code is available on Github (https://github.com/jennybigman/dsem_for_ebfm), which can guide others to use DSEM for these and other EBFM applications.

### Estimating ecosystem-level trends in the Chukchi Sea

#### Background

In subpolar ecosystems such as the Chukchi Sea, climate change is expected to cause a decrease in sea-ice extent and a change in the seasonal timing of the spring phytoplankton bloom (Fujiwara et al., 2016; Wang et al., 2012). This will likely cause a decrease in large ice algae that rapidly sink and support a rich community of benthic invertebrate consumers (Gradinger, 2009; Niemi et al., 2024). In turn, the benthic energetic pathway may instead be replaced by smaller phytoplankton supporting pelagic fish consumers (Lee et al., 2012). Such ecosystem changes may weaken the benthic pathway that supports clams, worms, crabs, and other invertebrates that are key prey species for marine mammals and are subsistence and food resources for many Arctic coastal communities. Ecosystem changes such as these can affect what is available to harvest, where resources are found, and seasonal harvest predictability (see Wise et al. 2025).

Composite indicators representing the total region-wide biomass of pelagic versus benthic consumers can be used to monitor an expected shift in “benthic-pelagic coupling”, and track shifts in the absolute values or relative composition of these communities. However, monitoring pelagic versus benthic biomass is difficult because these communities are typically sampled using different gears, and individual species within those communities will be sampled in some years and not others. Here, we demonstrate the use of DSEM to estimate time series of two composite variables, separately representing pelagic and benthic biomass. This analysis differs from Dynamic Factor Analysis (DFA), where multiple variables are shrunk towards one or more shared trends (Zuur et al., 2003). By contrast, Composite Factor Analysis (CFA) incorporates autocorrelation within each variable, allowing interpolation of missing years, but does not shrink time-series towards one another. Instead, it calculates composite factors as the weighted average of individual time-series (Buckland et al., 2005). Here, we use equal weighting when combining the log-transformed and centered values of each taxon classified as benthic or pelagic.

#### Hypothesis and inferential goal

We hypothesize that pelagic and benthic biomass will show divergent trends in the Chukchi Sea ecosystem, consistent with prior work suggesting a shift in benthic-pelagic coupling driven by declining sea ice and earlier phytoplankton blooms. The temporal dynamics of any single species or group over time would not adequately capture these community-level dynamics. Given this, our inferential goal is to estimate latent trends that represent changes in community-level, aggregated biomass. We are not interested in ascribing causality to environmental or other drivers, are not focused on statistical predictive performance, and as such, interpret only the trajectories of the trends rather than specific effects (path coefficients). We consider this a structural model rather than a structural causal model. DSEM offers a way to synthesize multiple, incompletely sampled time series into composite factors that can be tracked over time.

#### Data

To demonstrate CFA, we assemble a database of ecological samples from the Chukchi Sea for 2002-2020, collected using a combination of gears including (1) acoustic and midwater-trawl; (2) large mesh “otter” trawl; (3) small mesh “beam” trawl; (4) Niskin bottom water sampling; and (5) pelagic bongo net trawls. Species measurements are then aggregated taxonomically to develop 27 “taxa” (ranging from species to class; see Wise et al. 2025 for more detail). For each taxon, sampling data are fitted using a spatiotemporal generalized linear mixed model (GLMM) configured for index standardization. We specified a log-linked Tweedie distribution with a fixed effect for year, a spatial Gaussian Markov random field (GMRF), and a spatiotemporal GMRF that follows a random walk over time. For each taxon, we then predicted total biomass-density across the Chukchi Sea for any year with available samples and combined these 27 taxa into a time-series matrix. Taxa range widely in available data from two years of sampling (Alaska plaice) to 17 years of sampling (Crustacea) (Figure 1). We then specified a DAG for the CFA in which each taxon contributed to its corresponding composite group, benthic or pelagic, with arrows directly linking the taxon to its group (Figure 2).

**Figure 1.**
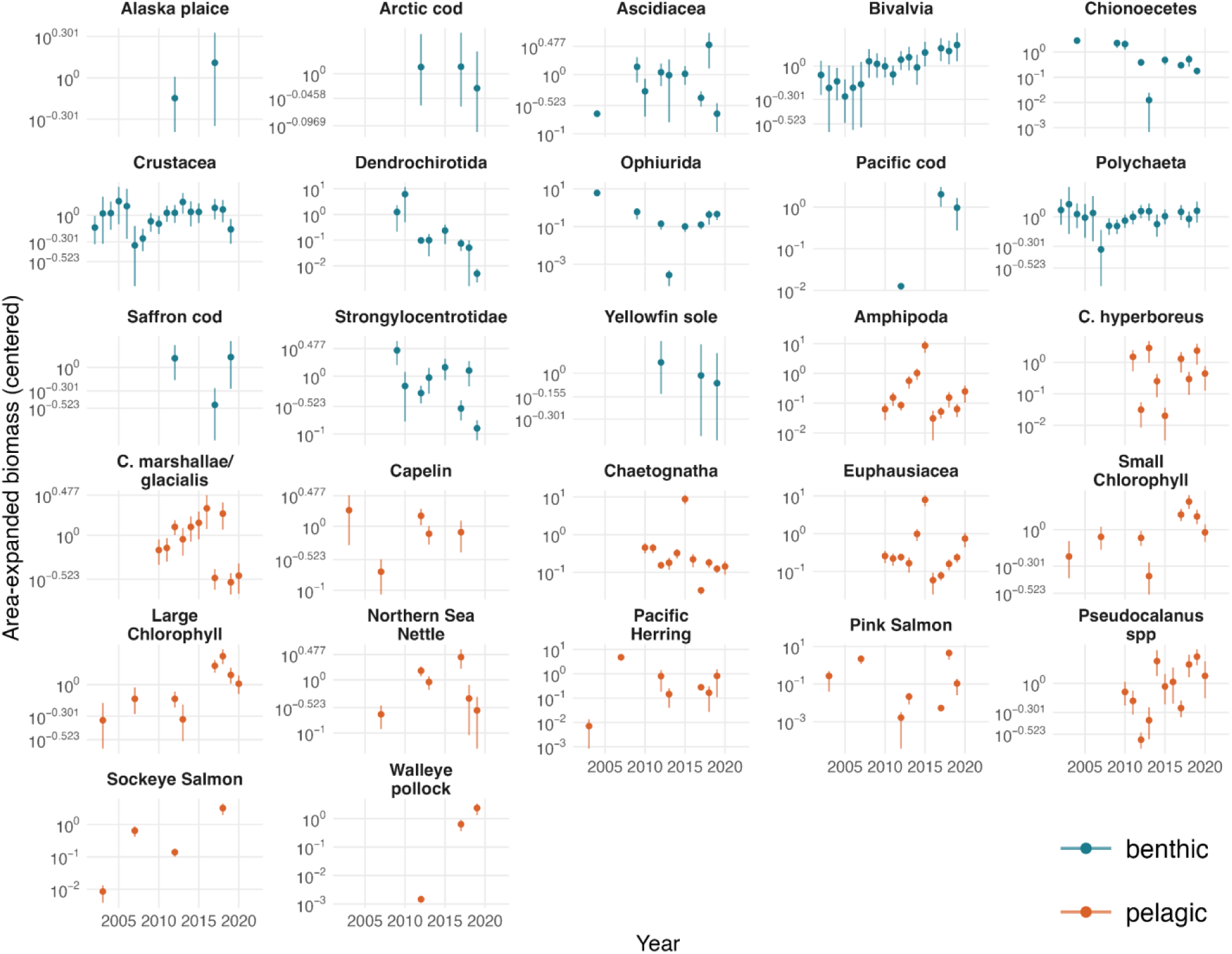
Relative index of area-expanded biomass and 95% confidence intervals (y-axis) from 2002-2020 (x-axis) in the Chukchi Sea for 27 taxa (panels) classified as primarily pelagic (blue) or benthic (orange), estimated by applying a spatiotemporal index-standardization model to data assembled across multiple gears. These model outputs are used as data in a subsequent DSEM model.

**Figure 2.**
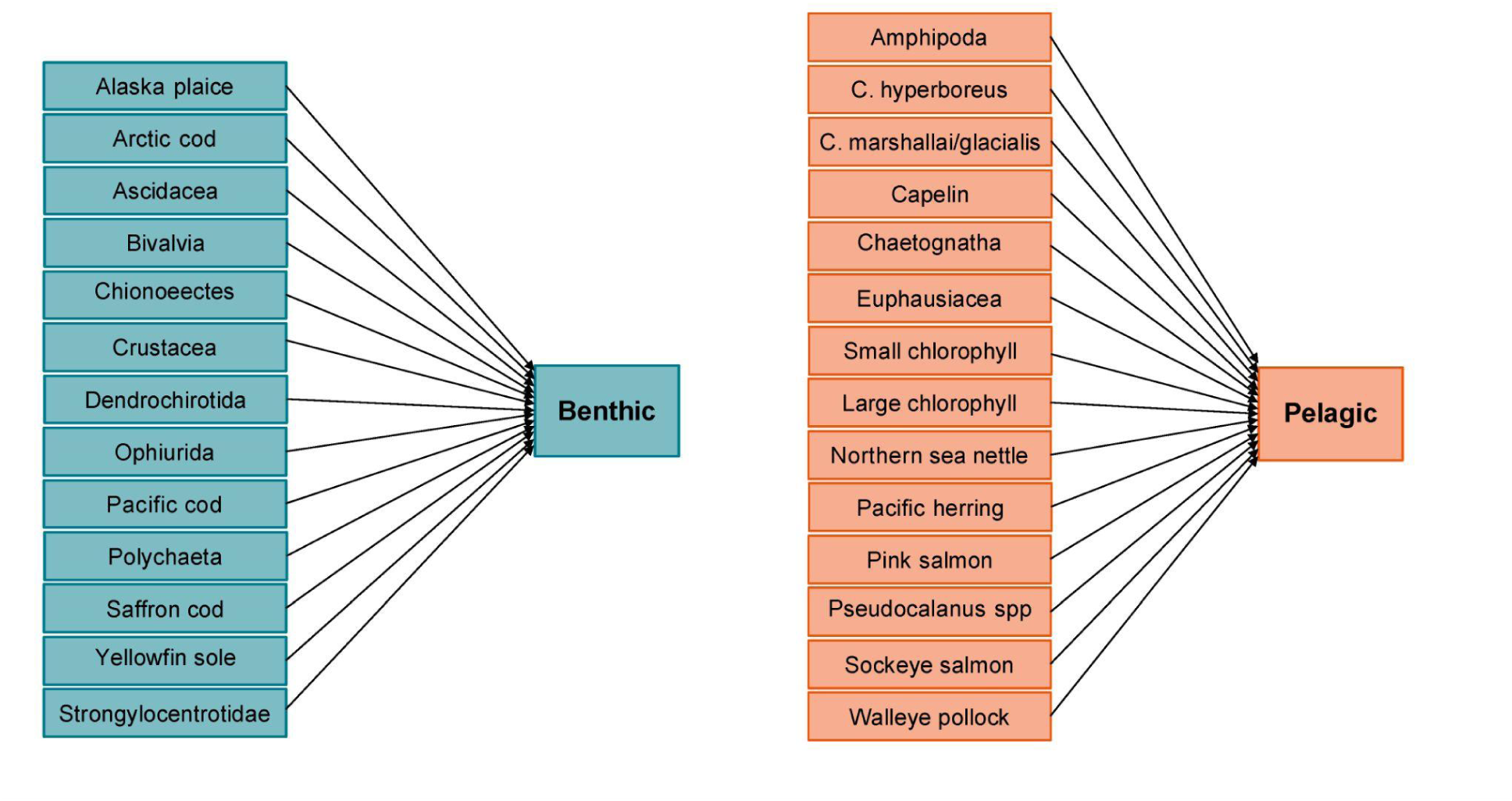
Directed acyclic graph representing the composite factor analysis (CFA) used in the first case study to estimate latent trends in aggregated benthic and pelagic biomass. In contrast to a conventional dynamic factor analysis (DFA), arrows originate from the individual taxon and point toward the corresponding composite group, indicating that the benthic and pelagic factors are constructed from their assigned observed time series. For DFA, the arrows would originate from the group (benthic or pelagic) and point towards each individual taxon.

#### Model

Using these time series and the DAG (Figure 2), we fit this DSEM model with the {dsem} package (version 2.0.1.9) in R v4.4.2 (Thorson et al., 2024; R core team 2025). The model estimates a single parameter (‘rhò) that defines the magnitude of first-order autocorrelation for all 27 taxa. We also fix 27 loadings parameters, where each loading has a value of one and points from a taxon to either benthic or pelagic composite variables. Finally, we estimate 27 parameters representing the conditional variance for each taxon. In total, the model estimates 28 fixed effects and then interpolates missing values for each taxon as well as the resulting value for the two composite variables.

## Results

Contrary to expectations, the composite indices showed little directional changes in the Chukchi Sea from 2002-2020 (Figure 3). Instead, the benthic index has multi-year cycles with peaks in 2004, 2009, and 2018 and a statistically significant low in 2013. Similarly, the pelagic index has peaks in 2007, 2015, and 2018, and a low in 2012. Confidence intervals are wide for most years, due to the infrequent sampling that is available in the Chukchi Sea for most species. We therefore conclude that (1) accurate detection of changes in benthic versus pelagic biomass would likely require more frequent biological sampling across ecosystem components, and (2) here, multi-year fluctuations are more supported than an overall directional trend for this system. However, wide confidence intervals around the composite trends limit our ability to draw strong conclusions from this analysis alone.

**Figure 3.**
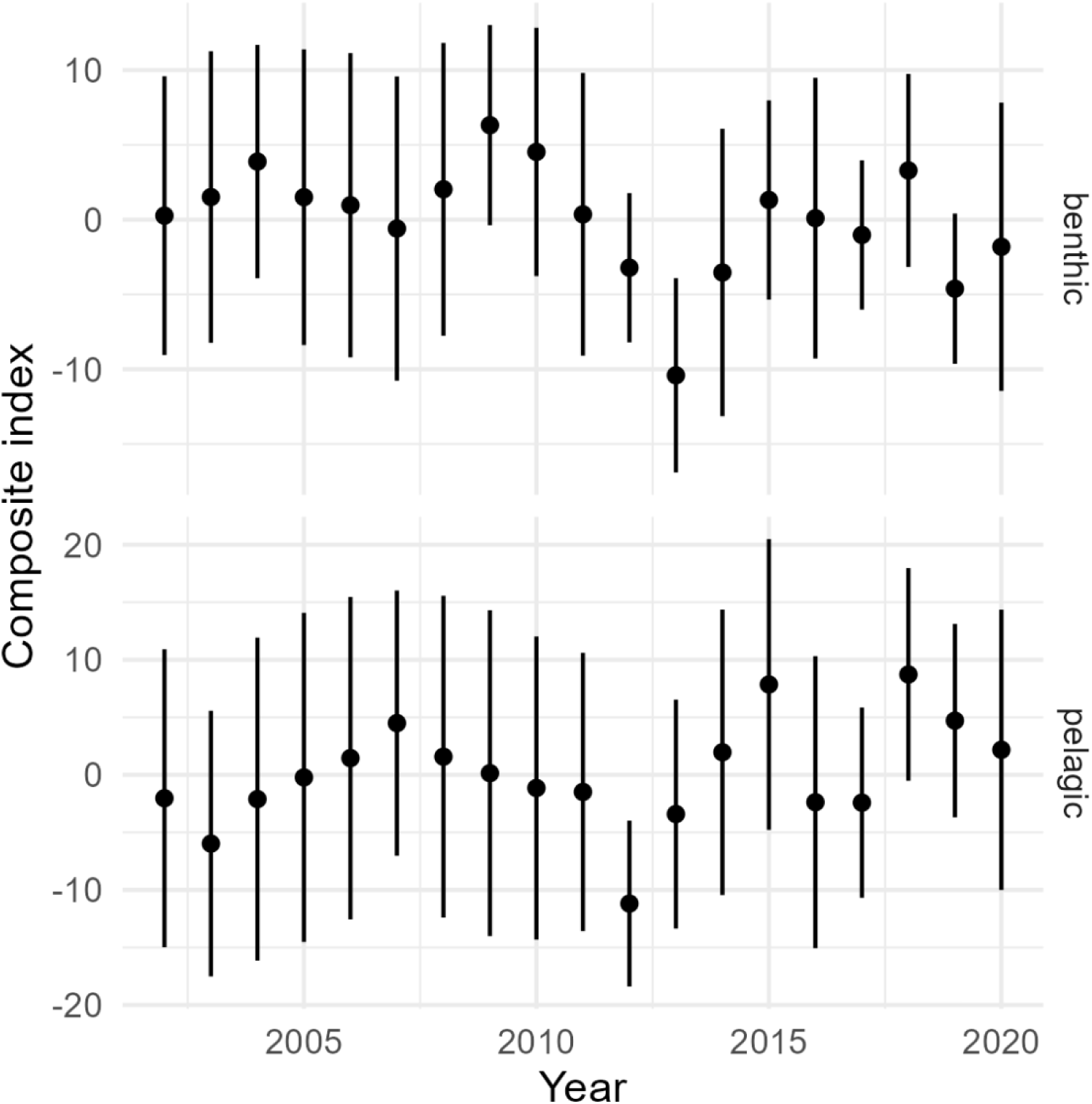
Composite indices (points) and 95% confidence intervals (vertical lines) for benthic (top panel) or pelagic (bottom panel) biomass (y-axis) for each year 2002-2020 (x-axis). Indices are calculated using a DSEM model that fits a first-order autoregressive process for log-biomass (i.e., a Gompertz model) for area-expanded relative biomass of each taxon (shown in Fig. 1), and jointly predicts composite indices as the sum of log-relative-biomass across benthic or pelagic taxa.

### Gulf of Alaska arrowtooth flounder recruitment

#### Background

In the Gulf of Alaska (GOA), arrowtooth flounder (*Atheresthes stomias*) is one of the most abundant groundfish species and has the highest market value and catch volume of any flatfish in the region (Spies et al., 2017; Shotwell et al., 2025). This species is an important predator of other groundfish, with trophic links to many other species in the system such as walleye pollock (Adams et al., 2022; Aydin et al., 2007; Barnes et al., 2020). As for many fishes, arrowtooth flounder recruitment is likely affected by many ecosystem factors both directly and indirectly, and is variable over time, with an apparent shift to lower recruitment beginning in 2006 (Doyle et al., 2018; Ward et al., 2024; Spies et al., 2017). There has also been a reduction in biomass of this species (along with other groundfishes) that preceded the recent marine heatwaves in the Gulf of Alaska, which may be linked to decreased recruitment (Spies et al., 2017; Shotwell et al., 2025). Due to the abundance and ecological importance of arrowtooth flounder, changes in the population dynamics of this species likely have a large effect on the ecosystem (Doyle et al., 2018; Shotwell et al., 2025; Spies et al., 2017).

#### Hypothesis and inferential goal

We hypothesize that arrowtooth flounder recruitment is governed by a combination of direct and indirect ecosystem linkages. Given this, our inferential goal is to develop a model that focuses on predictive (or statistical) performance and best predicts arrowtooth flounder recruitment rather than the interpretation of individual effects. We use ecological knowledge to develop a structural model but do not aim to test or advance causal understanding. Therefore, we are not concerned about potential confounding bias, inconsistency between the data and structural assumptions, or unexpected signs of effects between variables. DSEM offers a way to extend regression to include indirect effects and autocorrelated time series.

#### Data

Time series of ecosystem factors were developed as part of the ESP process, which is underway for arrowtooth flounder (Shotwell et al., 2025). The linkages included one outcome variable, recruitment, and 10 ecosystem variables: bottom temperature, predation, prey for age-1 recruits, sea surface temperature, larval abundance, prey for larvae, retention dynamics (two metrics), body condition of adult arrowtooth flounder, and abundance of young-of-the-year fish. Age-1 recruitment was taken from the latest reviewed stock assessment (Shotwell et al., 2025). All data were standardized (centered and scaled) for model fitting, and we assume that variables are measured without error (i.e., a process-error model). The data, their sources, and specified latent structures for model fitting are described in detail in Table S1.

#### Model

To develop a DAG to explain how ecosystem components may interact to affect arrowtooth flounder recruitment, we consulted the literature and the team of experts that contribute to the ESP for this stock, which consists of ecological subject matter experts, ecosystem and economic status report representatives, the stock assessment leads, and the ESP facilitator. We identified a total of 10 linkages that may directly and indirectly inform arrowtooth flounder recruitment (Table 1, Figure 4). In addition to documenting the hypothesized linkages, we report the expected sign of the relationship (e.g., positive or negative), the lag, the literature source that supports the linkage, and our confidence in the strength or importance of the linkage (based on discussion with subject-matter experts, relevance to the specific stock versus populations in other regions, directly observed versus modeled, and body of literature supporting link) (Table 1). Such documentation in a ‘mechanism table’ is an important step in the development of a graphical model, as the goal is to understand how different processes interact to affect an outcome of interest (here, recruitment; Grace and Irvine, 2020).

**Figure 4.**
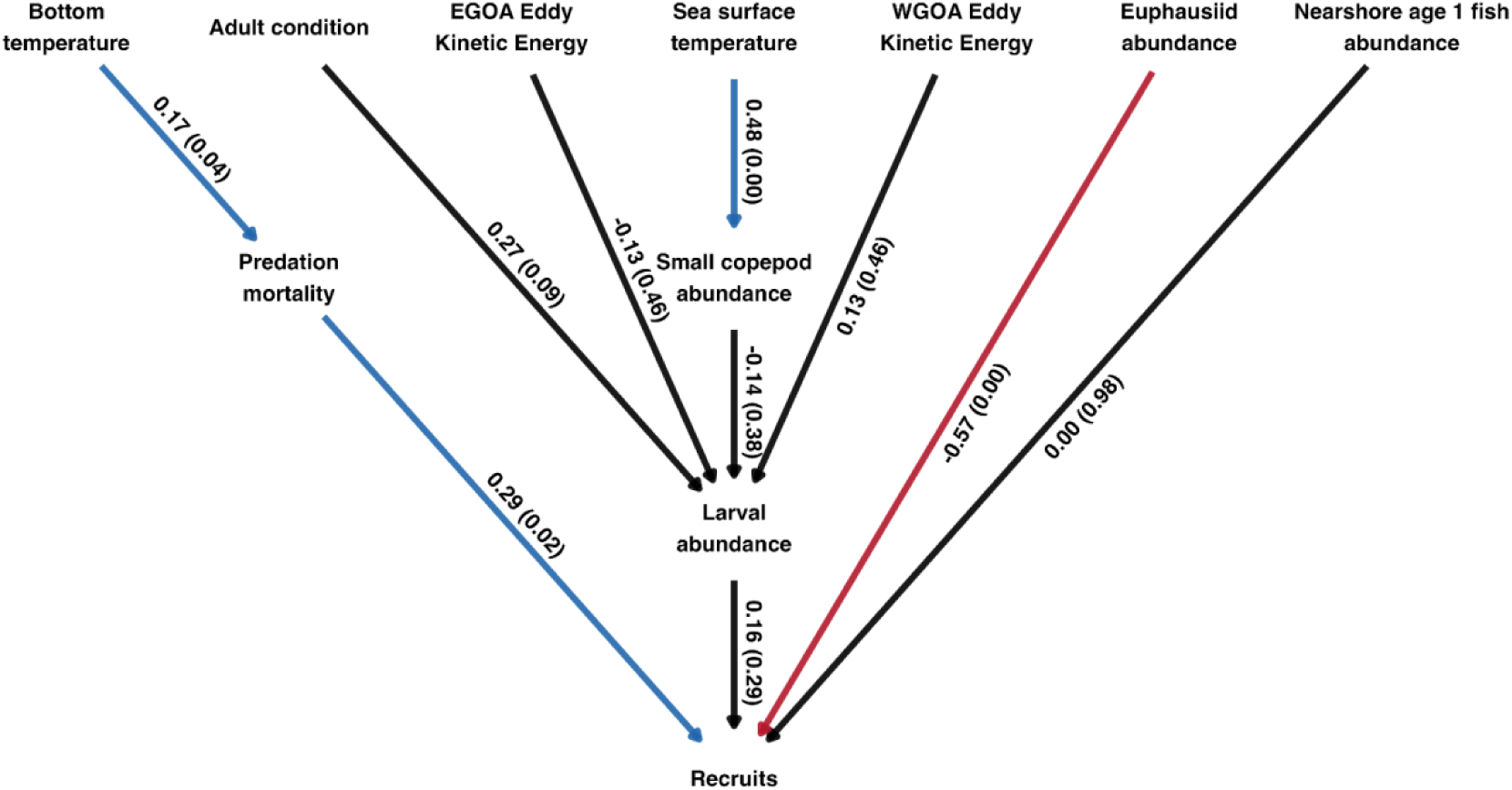
The causal model to explain arrowtooth flounder recruitment with estimated effects and p-values (in parentheses). The direction of the arrow indicates the hypothesized causal relationship and the color indicates the sign (red = negative, blue = positive, black is not significant). Lags and more detail about the variables can be found in Table 1.

**Table 1.**
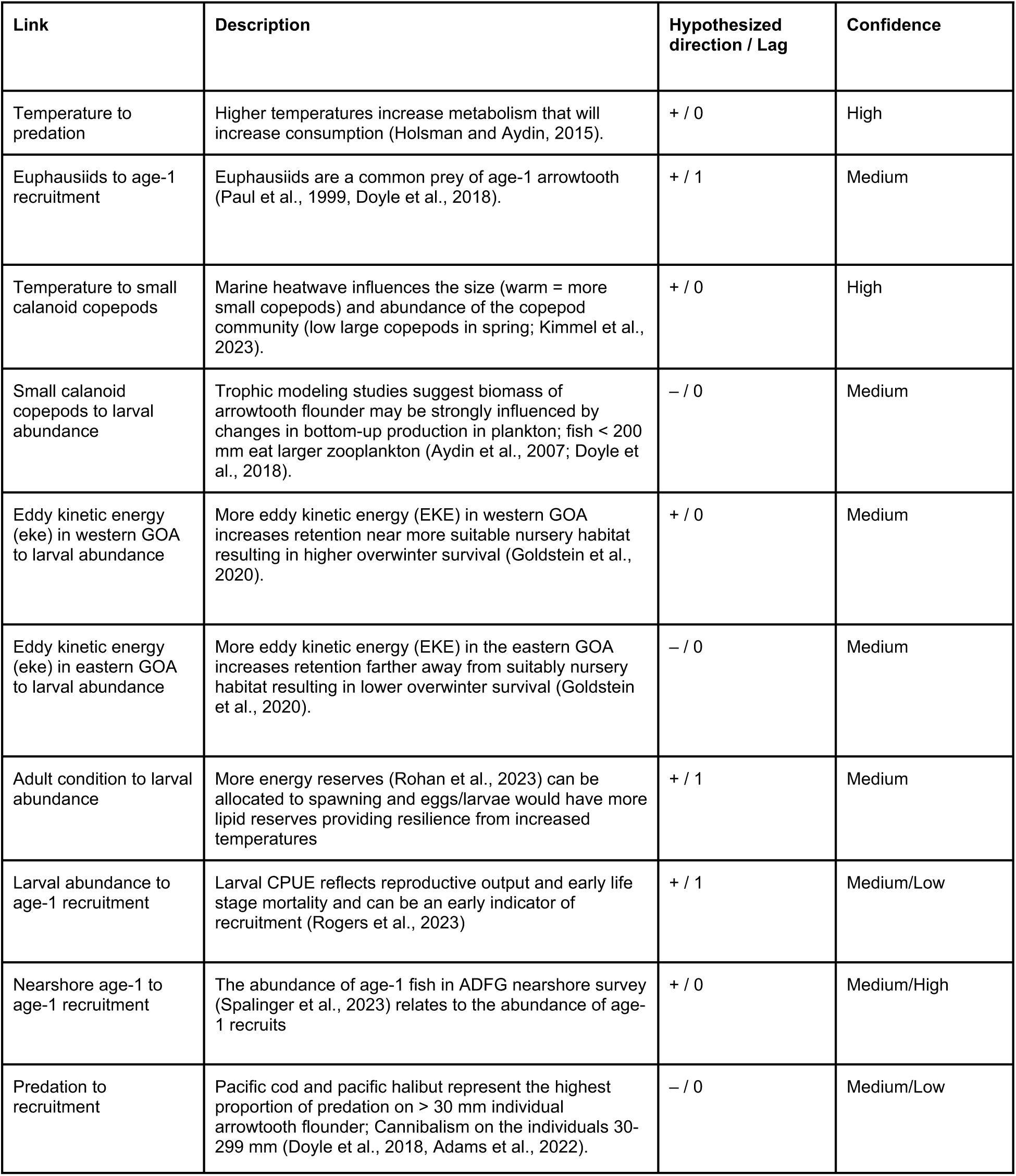
Description of direct linkages included in the directed acyclic graph (DAG) to explain recruitment of arrowtooth flounder in the Gulf of Alaska (GOA) sourced from expert opinion and the literature. We also include the hypothesized direction of the causal relationship, lag of effect, and expert confidence in the linkage.

| Link | Description | Hypothesized direction / Lag | Confidence |
| --- | --- | --- | --- |
| Temperature to predation | Higher temperatures increase metabolism that will increase consumption (Holsman and Aydin, 2015). | + / 0 | High |
| Euphausiids to age-1 recruitment | Euphausiids are a common prey of age-1 arrowtooth (Paul et al., 1999, Doyle et al., 2018). | + / 1 | Medium |
| Temperature to small calanoid copepods | Marine heatwave influences the size (warm = more small copepods) and abundance of the copepod community (low large copepods in spring; Kimmel et al., 2023). | + / 0 | High |
| Small calanoid copepods to larval abundance | Trophic modeling studies suggest biomass of arrowtooth flounder may be strongly influenced by changes in bottom-up production in plankton; fish < 200 mm eat larger zooplankton (Aydin et al., 2007; Doyle et al., 2018). | - / 0 | Medium |
| Eddy kinetic energy (eke) in western GOA to larval abundance | More eddy kinetic energy (EKE) in western GOA increases retention near more suitable nursery habitat resulting in higher overwinter survival (Goldstein et al., 2020). | + / 0 | Medium |
| Eddy kinetic energy (eke) in eastern GOA to larval abundance | More eddy kinetic energy (EKE) in the eastern GOA increases retention farther away from suitably nursery habitat resulting in lower overwinter survival (Goldstein et al., 2020). | - / 0 | Medium |
| Adult condition to larval abundance | More energy reserves (Rohan et al., 2023) can be allocated to spawning and eggs/larvae would have more lipid reserves providing resilience from increased temperatures | + / 1 | Medium |
| Larval abundance to age-1 recruitment | Larval CPUE reflects reproductive output and early life stage mortality and can be an early indicator of recruitment (Rogers et al., 2023) | + / 1 | Medium/Low |
| Nearshore age-1 to age-1 recruitment | The abundance of age-1 fish in ADFG nearshore survey (Spalinger et al., 2023) relates to the abundance of age-1 recruits | + / 0 | Medium/High |
| Predation to recruitment | Pacific cod and pacific halibut represent the highest proportion of predation on > 30 mm individual arrowtooth flounder; Cannibalism on the individuals 30-299 mm (Doyle et al., 2018, Adams et al., 2022). | - / 0 | Medium/Low |

Using the time series and the DAG (Figure 4), we fit a model using DSEM to compare how well our ‘structural model’ explains recruitment over time. For comparison, we fit three additional models using DSEM: one modeling recruitment as independent and identically distributed (iid), one modeling recruitment as a first order autoregressive (AR1) process, and one model that includes only variables hypothesized to directly relate to recruitment (larval abundance, abundance of young of the year fish the year prior, predation, and prey for age-1 recruits; ‘regression’). We ensured all models converged (all absolute marginal log-likelihood gradients were < 0.001 and the Hessian matrix of second derivatives of the marginal negative log-likelihood was positive-definite). We evaluated model fit using leave-one-out residuals and compared relative predictive performance for these three models using Akaike’s Information Criterion (AIC) (Figures S1-S4; Burnham & Anderson, 2002; Thorson et al., 2024). We estimated the total effect of each predictor on recruitment, which combines direct and indirect effects at a given lag and helps identify the variables with the greatest influence on recruitment (Table 2). We calculated total effects using the *total_effect* function in the {dsem} R package under a pulse-response formulation (Thorson et al., 2024). In this formulation, the function applies a one-unit change to a focal predictor at (t=0) and tracks how the effect propagates through the system over subsequent time steps (here, 0-3 lags). Total effects incorporate all direct and indirect pathways connecting a predictor to recruits, including temporal propagation through autoregressive relationships. We summarized each predictor–recruit relationship using the total effect with the greatest absolute magnitude across the subsequent lags (Table 2).

**Table 2.** Value of the total effect (maximum across lags) on recruitment in the structural model. Abbreviations: SST = sea surface temperature, copepods = small copepod abundance, larvae = larval abundance, age1 = recruits (abundance of age 1 fish), eke_east = eastern Gulf of Alaska eddy kinetic energy, eke_west = western Gulf of Alaska (GOA) eddy kinetic energy, cond = adult condition, nearshore age1 = nearshore age 1 fish abundance, bt = bottom temperature, pred = predation mortality, euph = euphausiids. Lag noted in the table is the lag associated with the maximum total effect.

| Pathway | Total effect (maximum<br>across lags) | Lag |
| --- | --- | --- |
| sst → copepods → larvae → age1 | -0.01 | 1 |
| copepods → larvae → age1 | -0.02 | 1 |
| eke_east → larvae → age1 | -0.02 | 1 |
| eke_west → larvae → age1 | 0.02 | 1 |
| cond → larvae → age1 | 0.04 | 2 |
| larvae → age1 | 0.16 | 1 |
| nearshore_age1 → age1 | 0.00 | 0 |
| bt → pred → age1 | 0.05 | 0 |
| pred → age1 | 0.29 | 0 |
| euph → age1 | -0.57 | 1 |

#### Results

The DAG for the structural model represented hypothesized linkages between environmental and ecosystem factors and Gulf of Alaska arrowtooth flounder recruitment (Table 1, Figure 4). We found that all models converged and had reasonable residuals (Fig. S1-S4). Although the structural model had the best predictive performance as measured by AIC, the structural and regression models both reduced unexplained variation substantially (82% and 83%, respectively; Table 3). In the structural model, we found strong evidence that several variables were associated with recruitment, such as prey availability (euphausiid abundance, maximum total effect = -0.57 and predation mortality (maximum total effect = 0.29), but that some hypothesized direct and indirect links were insignificant (Table 2, Figure 4). The interpretation of the total effects is the same as in a traditional regression, namely that we have quantified statistical associations among variables given a hypothesized model and identified which variables are useful for prediction, but not counterfactual (causal) claims as that was not the inferential goal and appropriate scientific validation was not done (see subsequent case study for scientific validation steps). The structural model also provided estimates (and associated uncertainty bounds) for ecosystem indicators in years without direct measurements (Figure 5), allowing us to better describe ecosystem status despite inconsistent field sampling among variables. Overall, our structural model suggested that many interacting environmental and ecosystem variables can be used to explain substantial variation in arrowtooth flounder recruitment, and this can be used to project recruitment and improve proposed catch in fisheries management systems (Champagnat et al., 2026).

**Figure 5:**
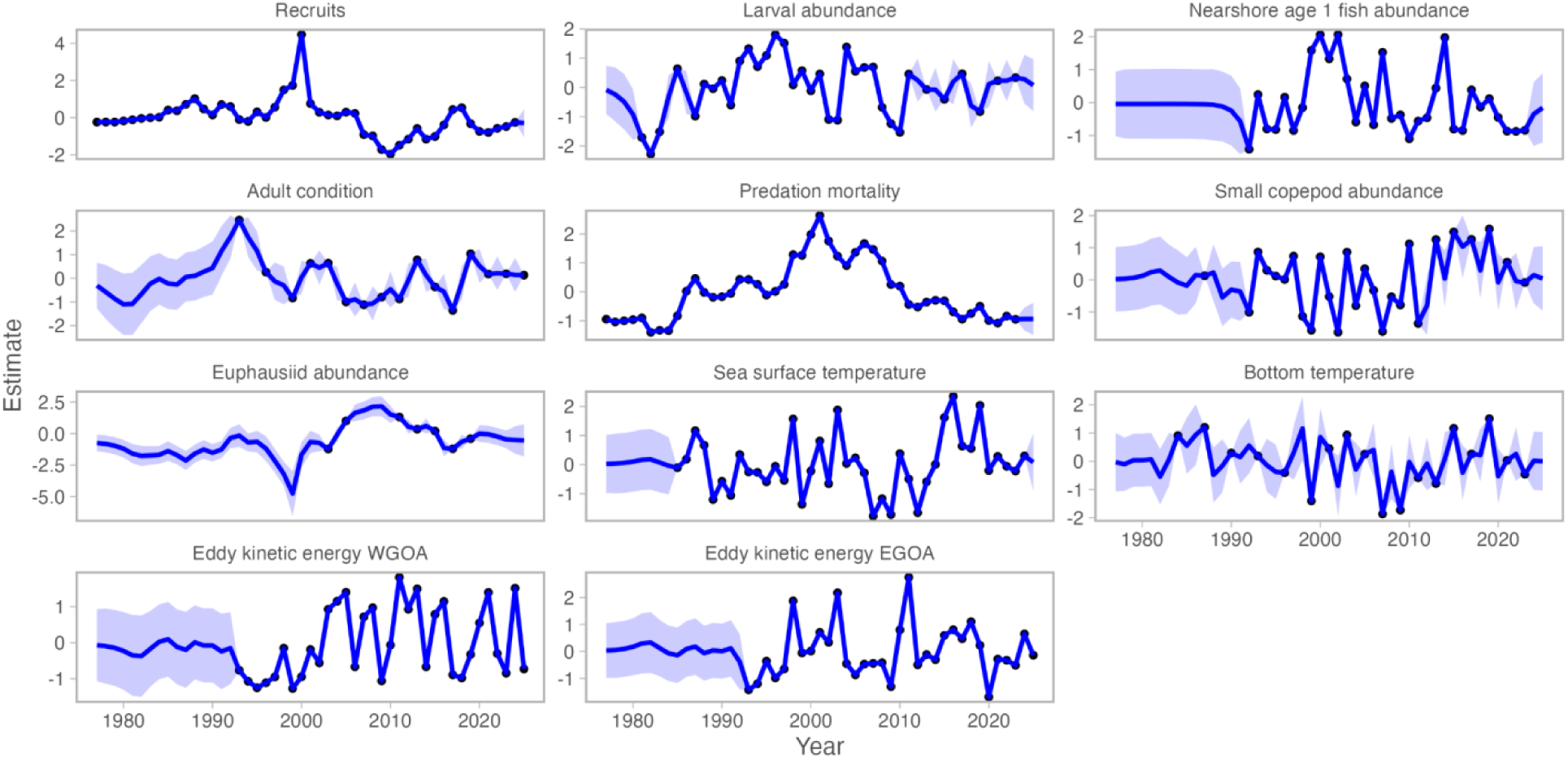
Time series of scaled recruitment (outcome) and covariates estimated using a dynamic structural equation model (DSEM) fitted to the causal model in Figure 2.

**Table 3.** Model selection results for the case study of Arrowtooth flounder recruitment. The structural model (represented by the DAG in Fig. 3) had better predictive accuracy as measured by AIC compared to modeling recruitment a first order autoregressive process (ar1), as independent and identically distributed (iid) or modeling recruitment, or as a function of the direct links in the DAG (regression, see text). ***σ_R, unexplained_*** is the value of the standard deviation of the response variable (recruits), indicating variability in the response that is unexplained by the model, and the reduction in this unexplained variation from the iid model.

| Model | AIC | $\Delta\text{AIC}$ | $\sigma_{R, \text{unexplained}}$ | % reduction in unexplained variation |
| --- | --- | --- | --- | --- |
| structural | 852.5 | 0 | 0.420 | 82% |
| regression | 855.3 | 2.7 | 0.416 | 83% |
| ar1 | 858.7 | 6.2 | 0.716 | 49% |
| iid | 888.4 | 35.9 | 1.000 | 0% |

### Understanding drivers of fish productivity on the U.S. Northeast Shelf

#### Background

Understanding how changes in environmental and ecosystem variables affect the productivity of fish stocks is central to ecosystem-based fisheries management (Link, 2002; Pikitch et al., 2004). For managed fishes on the U.S. Northeast continental shelf, there have been a series of studies that have explored how prey availability and demography (e.g., the abundance and condition of spawners) affect the productivity of fishes within and across fish stocks (Friedland et al., 2023; Perretti et al., 2017; Pershing et al., 2005). Many of these studies have focused on the relationship between the abundance of zooplankton and young fish, how biomass and condition of spawners may affect young fish abundance, or how demography and prey availability may interact to affect the fish abundance (Fogarty & Cohen, 1991; Friedland et al., 2023; Perretti et al., 2017). Using causal inference to compare support for different hypotheses offers an opportunity to re-examine these mechanistic relationships that were previously identified with non-causal techniques.

#### Hypothesis and inferential goal

Here, we build on this body of work and evaluate support for three causal hypotheses regarding the abundance of young fish on the U.S. Northeast Shelf (Figure 6). The first hypothesis suggests that the abundance of young fish is driven by the abundance of copepods (‘prey availability’). The second hypothesis posits that the abundance of young fish is driven directly by the biomass and condition of spawners, as well as the density-dependent effect of biomass on condition (i.e., the stock-recruit relationship may be influenced by condition, ‘demography’). Finally, the third hypothesis suggests that both prey availability and demography act together to affect the abundance of young fish (‘prey availability + demography’). Through this case study, we demonstrate how to develop, compare, and validate graphical models that reflect distinct causal hypotheses. Our inferential goal is to compare support among these three mechanistic hypotheses for as many populations of fish in the region as possible. Unlike previous case studies, here, we are interested in causally interpreting the effects and identifying the most supported causal structure. This requires additional scientific validation steps, such as ensuring the effect signs are biologically plausible and the DAG is consistent with the data. In this case study, we will demonstrate the steps to scientifically validate models using DSEM.

**Figure 6.**
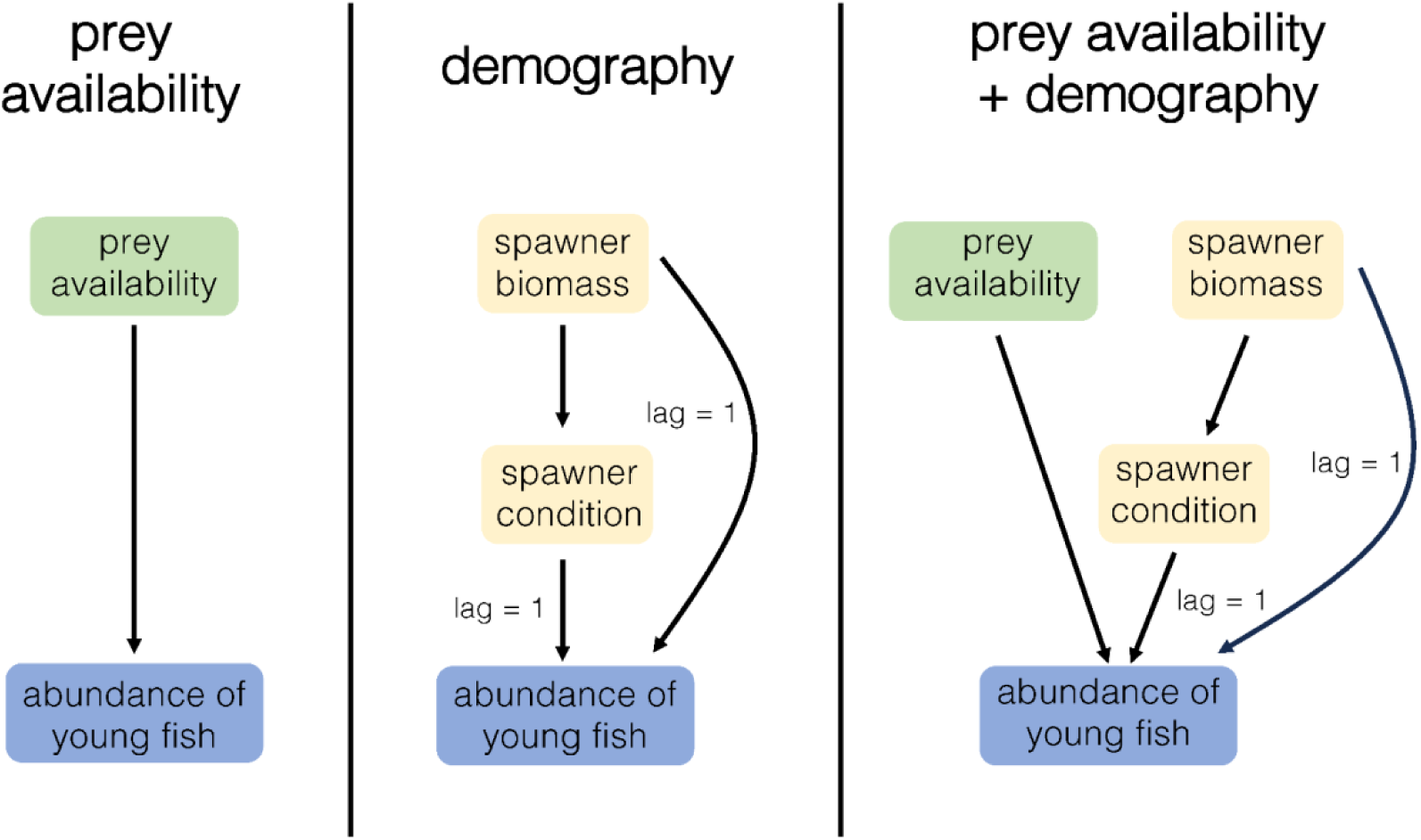
The three competing hypotheses explaining the abundance of young fish on the U.S. Northeast Shelf. In the second and third panel (demography and prey availability + demography), a 1-year lag is included between spawner biomass and the abundance of young fish and spawner condition and abundance of young fish.

#### Data

Time series of physical and biological variables on the Northeast Shelf of the U.S. were taken from the {ecodata} R package (Beltz et al., 2025) and the NEFSC bottom trawl survey, as described in Azarovitz (1981) and Smith (2002). We included data for all species collected in the survey that were subsequently aged (n = 25), and species were grouped into populations based on their corresponding ‘Ecological Production Unit’ (EPU): Gulf of Maine (GOM), Mid Atlantic Bight (MAB), and Georges Bank (GB) (see Perretti et al., 2017). The response variable, ‘abundance of young fish,’ is the abundance of age 0 and age 1 fish of a given species in a given EPU and the predictor variables were a metric of prey availability—the annual abundance anomaly of small copepods (‘small copepod anomaly’), annual mean condition (relative ‘condition’ of the 74 populations), and the biomass of large fish (age 2 and older) from the survey (‘large fish biomass’).

#### Model

We first developed three DAGs that reflected our causal hypotheses (prey availability, demography, or prey availability + demography) regarding fish productivity (Figure 6). Then, for each population (a stock in a given EPU), we fit a DSEM model for each of these three hypotheses. We used a directional separation test (‘d-separation’) for all three DAGs for all populations to check whether our DAG was consistent with the observed dependency structure in the data (Arif & MacNeil, 2023; Shipley, 2000). The d-separation test is currently in development for DSEM but provides a useful tool for examining implied conditional independencies and evaluating whether the hypothesized causal assumptions encoded in the DAG are supported by the data. As such, we use it here to illustrate the utility and workflow. In the {dsem} package, a d-separation test is implemented in the *test_dsep* function, where all conditional independencies are sequentially tested and then combined in a single omnibus test (Thorson et al., 2025).

Our model comparison proceeded as follows: (1) models were excluded if they failed to converge according to the same criteria as the first case study or if the d-separation test yielded an omnibus p-value (p-value for the entire DAG) of < 0.1, indicating poor fit of the overall DAG (and thus corresponding hypotheses); and (2) when two or more models were retained for a given population, the models were compared using AIC, whereas a single retained model for a population was interpreted as the most supported hypothesis among those evaluated. Because separate DSEM models were fitted to 74 populations, the large number of estimated pathways increases the likelihood of Type I error; therefore, in our results section, we emphasize effect size, direction, and consistency across all populations rather than over-interpreting isolated effects and their p-values. Total effects were quantified as in the previous case study using the *total_effect* function in the {dsem} R package using a pulse-response formulation (Thorson et al., 2024).

### Results

Out of a total of 74 populations, models for 36 populations were retained following basic model checking and d-separation tests (Table 4). Whether prey availability (zooplankton abundance), demography (biomass and condition of spawners), or both prey availability and demography drove the abundance of young fish differed by population. For 23 of 36 populations, one causal hypothesis had more support over the other two based on convergence, d-separation tests, or AIC (Table 4). For most of these (17 of 23), prey availability alone was the causal driver of the abundance of young fish. Demography drove the abundance of young fish for four populations and both prey availability and demography drove the abundance of young fish for two populations (Table 4, Figure 7). Equal support for more than one causal hypothesis driving the abundance of young fish was evident for the remaining 13 populations: five populations had equal support for all three hypotheses, four had equal support for both demography and both prey availability and demography, three had equal support for prey availability and demography, and one had equal support for prey availability and both prey availability and demography (Table 4). For the 15 species with multiple populations, four showed support for the same hypotheses (prey availability for Witch flounder in GOM and MAB, for Goosefish in GB and GOM, and for Butterfish in GOM and MAB). The most supported model for the remaining species with multiple populations was variable, as was the most commonly supported model in a given EPU. As such there does not appear to be a pattern by species or region (Table 4).

**Figure 7.**
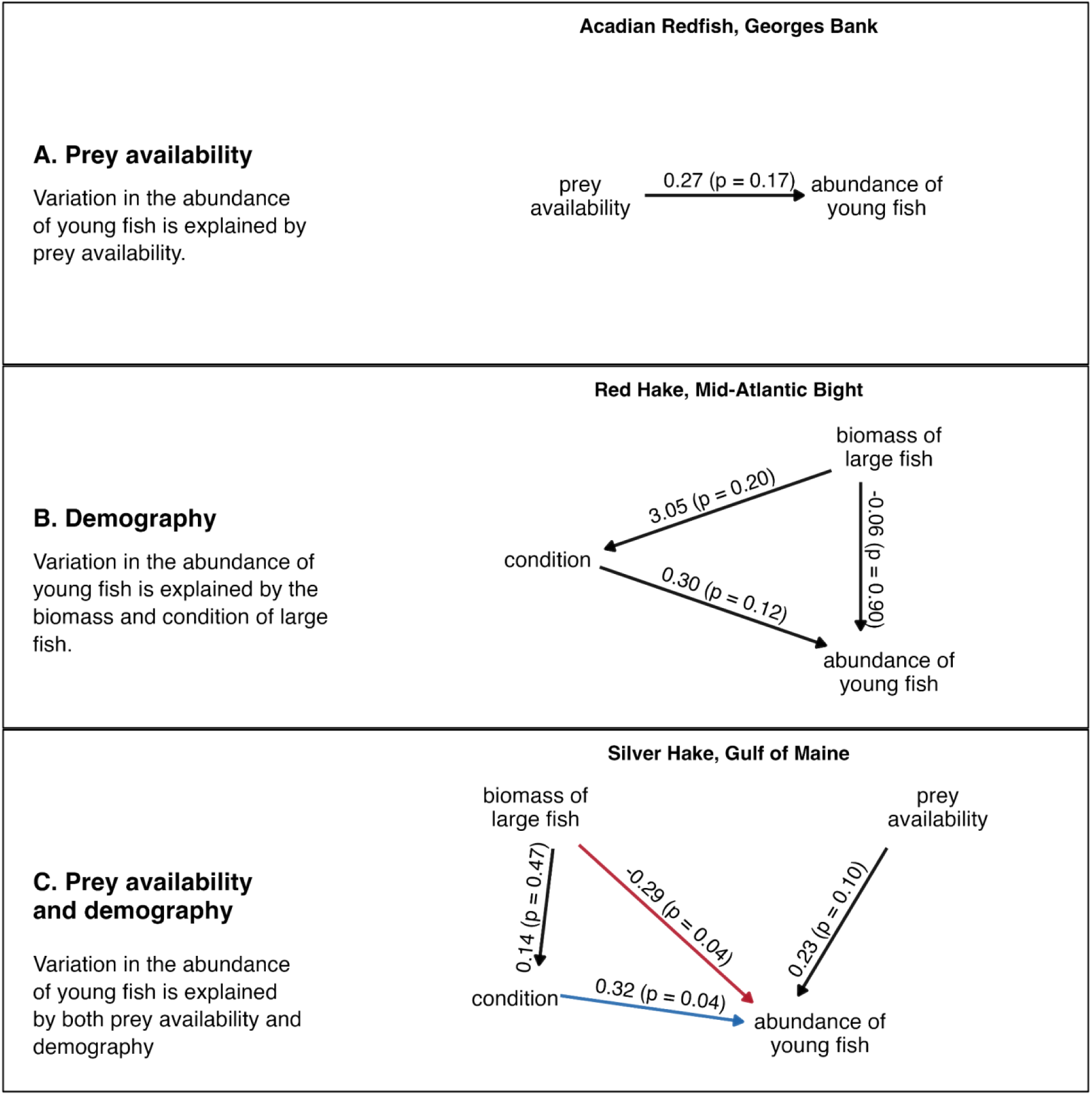
Causal hypotheses explaining variation in abundance of young fish for three (randomly selected) representative stocks on the U.S. Northeast Shelf, with each stock illustrating a different best-supported hypothesis (see text, Table 4). Numbers indicate the estimated effects and p-values (in parentheses). The direction of the arrow indicates the hypothesized causal relationship, and the color indicates the sign (red = negative, blue = positive, black = not significant). A 1-year lag was included for the link between spawner biomass and the abundance of young fish and the link between spawner condition and abundance of young fish.

**Table 4.**
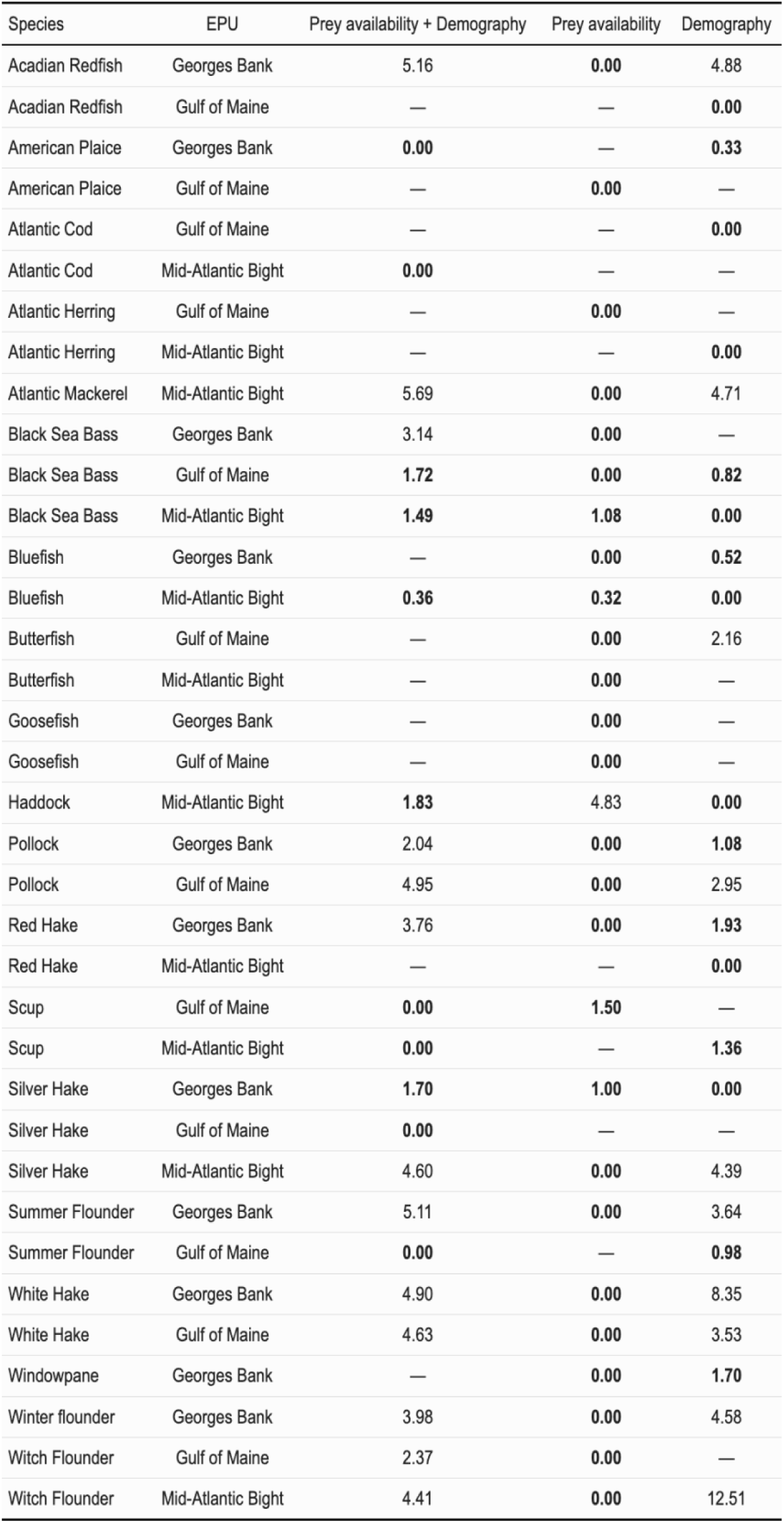
Delta (Δ) AICs of the three models specifying different causal drivers of the abundance of young fish on the U.S. Northeast Shelf for the 36 populations that were retained following convergence and d-separation tests (see text for more details). An emdash (–) indicates that the model did not converge or failed a d-separation test (omnibus p-value < 0.1, see text for more details).

Effect sizes, signs, and significance varied across the 17 populations that had support for prey availability driving the abundance of young fish (effect sizes ranged from -1.00 for MAB Witch flounder to 0.55 for GOM Goosefish, Table 5, Figure 7, S5). Most (11/17) of these had a non-zero, nonsignificant, positive effect of small copepod anomaly on the abundance of age 1 and 0 fish and the remaining populations had a negative effect (Table 5, Figure 7, S5). For the four populations for which demography was the most supported hypothesis, no effect was significant, the total effect of biomass on the abundance of young fish (direct and indirect through condition) was positive for all (ranged from 0.19 to 0.85), and the effect of adult condition on the abundance of young fish was positive for two populations and negative for the other two (ranged from -0.02 to 0.86) (Table 6, Figure 7, S5). Two populations–MAB Atlantic cod and GOM Silver hake– showed clear support for both prey availability and demography over either one of these alone. The effect of adult condition and the effect of small copepod anomaly on the abundance of young fish were positive for both populations (Table 7, S5). The total effect of biomass of large fish on the abundance of young fish differed in sign and magnitude between the two populations: for GOM Silver hake, a greater biomass reduced abundance of young fish but for MAB Atlantic cod, a greater biomass increased abundance of young fish (Table 7, S5).

**Table 5.** Value of the (maximum) total effect across lags on the abundance of young fish for the populations for which prey availability was the most supported model. Abbreviations: smcop = abundance anomaly of small copepods (see text), recruits = abundance of age 0 and 1 fish. Lag noted in the table is the lag associated with the maximum total effect.

| Population | Pathway(s) | Total effect (maximum across lags) | Lag |
| --- | --- | --- | --- |
| GB Acadian Redfish | smcop → recruits | 0.27 | 0 |
| GB Black Sea Bass | smcop → recruits | -0.41 | 0 |
| GB Goosefish | smcop → recruits | 0.00 | 0 |
| GB Summer Flounder | smcop → recruits | 0.14 | 0 |
| GB White Hake | smcop → recruits | -0.37 | 0 |
| GB Winter flounder | smcop → recruits | 0.37 | 0 |
| GOM American Plaice | smcop → recruits | 0.20 | 0 |
| GOM Atlantic Herring | smcop → recruits | -0.22 | 0 |
| GOM Butterfish | smcop → recruits | -0.07 | 0 |
| GOM Goosefish | smcop → recruits | 0.55 | 0 |
| GOM Pollock | smcop → recruits | 0.04 | 0 |
| GOM White Hake | smcop → recruits | 0.20 | 0 |
| GOM Witch Flounder | smcop → recruits | 0.19 | 0 |
| MAB Atlantic Mackerel | smcop → recruits | 0.15 | 0 |
| MAB Butterfish | smcop → recruits | 0.06 | 0 |
| MAB Silver Hake | smcop → recruits | -0.18 | 0 |
| MAB Witch Flounder | smcop → recruits | -1.00 | 0 |

**Table 6.**
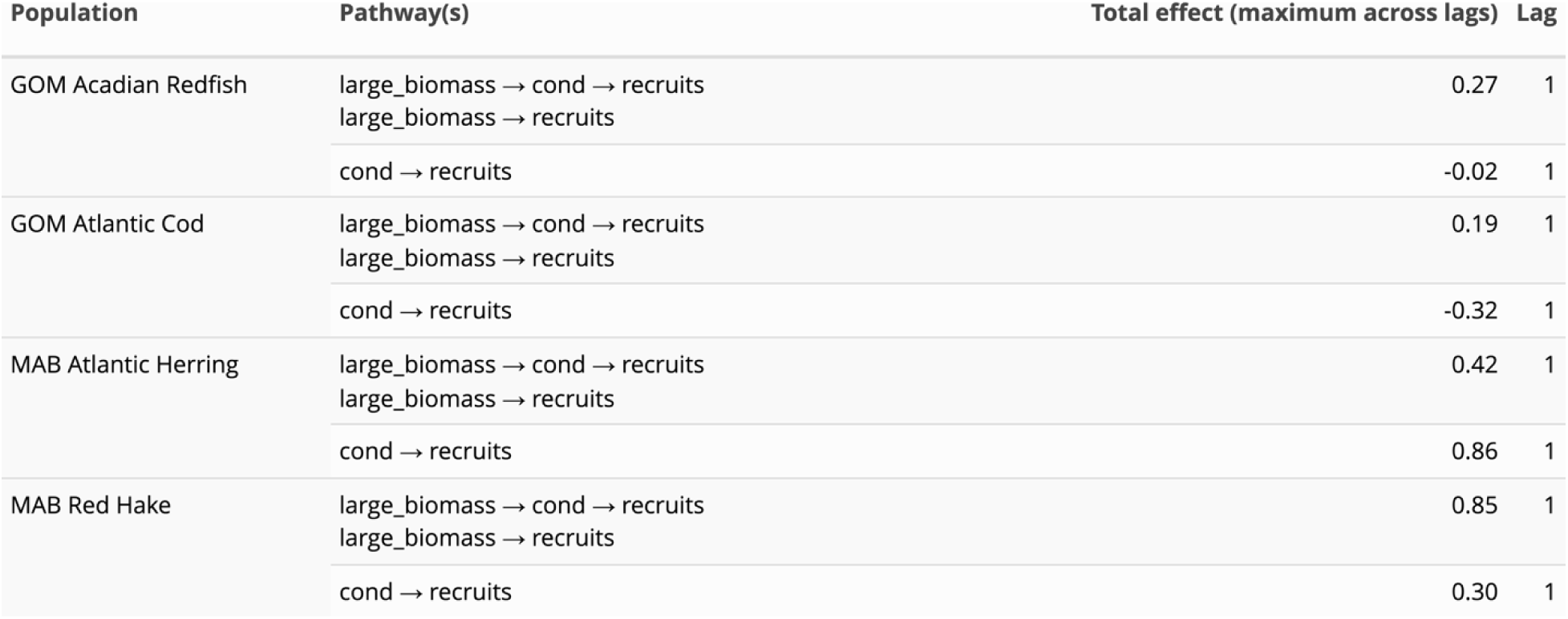
Value of the (maximum) total effect across lags on the abundance of young fish for the four populations for which demography was the most supported model. Abbreviations: large_biomass = biomass of age 2+ fish, cond = adult condition, recruits = abundance of age 0 and 1 fish. Lag noted in the table is the lag associated with the maximum total effect.

| Population | Pathway(s) | Total effect (maximum across lags) | Lag |
| --- | --- | --- | --- |
| GOM Acadian Redfish | large_biomass → cond → recruits | 0.27 | 1 |
|  | large_biomass → recruits |  |  |
|  | cond → recruits | -0.02 | 1 |
| GOM Atlantic Cod | large_biomass → cond → recruits | 0.19 | 1 |
|  | large_biomass → recruits |  |  |
|  | cond → recruits | -0.32 | 1 |
| MAB Atlantic Herring | large_biomass → cond → recruits | 0.42 | 1 |
|  | large_biomass → recruits |  |  |
|  | cond → recruits | 0.86 | 1 |
| MAB Red Hake | large_biomass → cond → recruits | 0.85 | 1 |
|  | large_biomass → recruits |  |  |
|  | cond → recruits | 0.30 | 1 |

**Table 7.** Value of the (maximum) total effect across lags on the abundance of young fish for the two populations for which prey availability and demography was the most supported model. Abbreviations: large_biomass = biomass of age 2+ fish, cond = adult condition, recruits = abundance of age 0 and 1 fish, and smcop = small copepod anomaly. Lag noted in the table is the lag associated with the maximum total effect.

| Population | Pathway(s) | Total effect (maximum across lags) | Lag |
| --- | --- | --- | --- |
| GOM Silver Hake | large_biomass → cond → recruits | -0.25 | 1 |
|  | large_biomass → recruits |  |  |
|  | cond → recruits | 0.32 | 1 |
|  | smcop → recruits | 0.23 | 0 |
| MAB Atlantic Cod | large_biomass → cond → recruits | 0.13 | 1 |
|  | large_biomass → recruits |  |  |
|  | cond → recruits | 0.94 | 1 |
|  | smcop → recruits | 0.07 | 0 |

## Discussion

Structural equation and structural causal models with DSEM allow for the integration of observational data, expert knowledge, and mechanistic hypotheses within a unified framework, while accommodating missing data and simultaneous and lagged interactions. Thus, DSEM offers a powerful tool for understanding the processes that govern complex ecosystems and advancing EBFM efforts. Collectively, the three case studies—estimating ecosystem-level latent trends, prediction of arrowtooth flounder recruitment, and comparing causal hypotheses governing drivers of fish productivity—demonstrate a few of the analyses that can be implemented in DSEM and highlight the flexibility of the framework for quantifying complex ecological interactions. Further, we provide examples for tailoring a SEM or SCM to a specific goal and synthesize this information into a workflow that broadly guides analysts in using DSEM for the different inferential goals explored in the study (Figure 8). In doing so, we address a key barrier to EBFM: the lack of a scientific and statistical framework capable of mechanistically describing ecosystem dynamics, accounting for correlated, indirect, and direct effects, and using observational data to identify causal relationships underlying ecosystem and population change.

**Figure 8.**
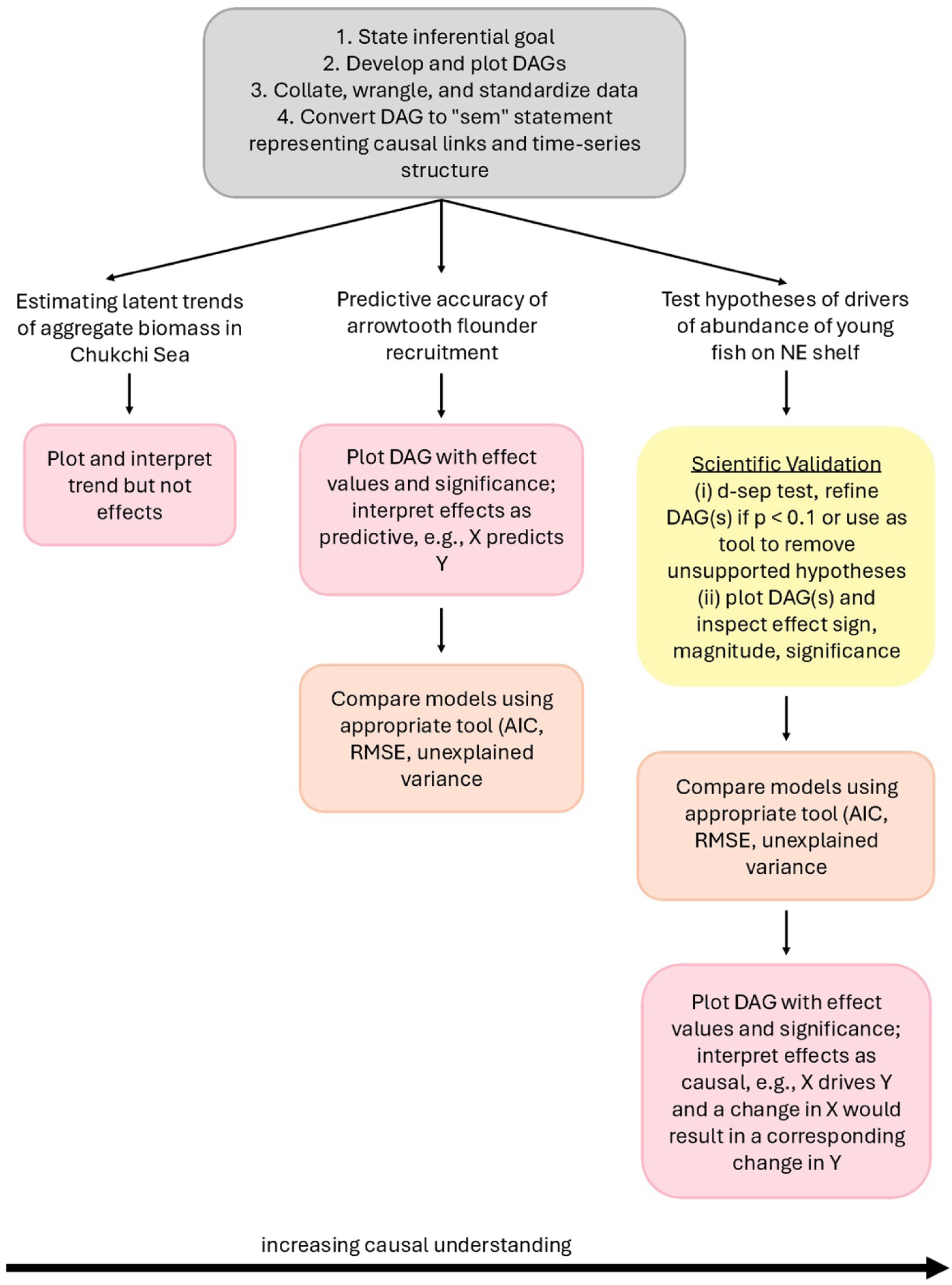
A workflow to help users navigate the different uses and applications of {dsem} for estimation, description, and causal inference. Gray boxes are steps common to all three case studies (and inferential goals) in this study, pink boxes indicate interpretation steps, orange boxes indicate model selection and comparison steps, and the yellow box indicates the steps that comprise scientific validation, which helps build causal understanding.

Our three case studies highlight how DSEM can provide information that is needed to support EBFM. For example, the first case study used DSEM to estimate broad-scale, ecosystem-level trends in biomass using composite indices. This approach has been used in monitoring biodiversity to combine trends of multiple groups in a standardized way and allow identification of change points (Buckland et al., 2005). Wise et al. (2025) compiled the ecosystem sampling data for the Chukchi Sea and then documented biomass changes for each species individually. However, ecosystem assessment often seeks to answer questions about species complexes and communities rather than individual species (Gove et al., 2022; Siddon, 2024). Analysts often accomplish this using dynamic factor analysis (DFA), in which latent factors are estimated as shared trends that explain variation among species-specific time series through species-specific loadings (i.e., factor → species; Zuur et al., 2003). By contrast, we used composite factor analysis, in which the shared trend is estimated as a composite of the species-specific time series, so that the observed time series informs the factor rather than being modeled as responses to it (i.e., species → factor). This demonstration is intended both (1) to estimate ecosystem indicators that avoid sharing information among species and (2) to demonstrate the flexibility of DSEM for developing ecosystem-level indicators.

Our second case study on arrowtooth flounder recruitment demonstrates how to incorporate both direct and indirect effects of ecosystem variables to increase predictive accuracy of recruitment. Indeed, the structural model–with both indirect and direct effects of ecosystem variables–had better predictive performance compared to modeling recruitment as independent and identically distributed, as a first order autoregressive process, or simply as a regression that included covariates hypothesized to have a direct effect on recruitment. DSEM also allows for the inclusion of unmeasured variables and the identification of confounding variables, or those that are mechanistically linked to both a predictor and the outcome and can explicitly accommodate lagged effects (in addition to simultaneous effects; Thorson et al., 2024). Lagged effects are common as ecosystems often do not immediately respond to drivers; for example, we hypothesized that the condition of adults affected larval abundance in the following year, larval abundance affected recruitment in the following year, and euphausiid abundance affected recruitment in the following year. Overall, this case study demonstrated DSEM’s great potential in supporting EBFM as understanding and quantifying the interrelationships among ecosystem and environmental variables and population dynamics is one major barrier to implementation (Frid et al., 2006). Specifically, our results give context to which measured variables can be an early warning sign for poor recruitment as part of the risk table approach (Dorn & Zador, 2020). To further support EBFM, the arrowtooth flounder DSEM model could be embedded in the stock assessment model to improve short-term forecasting of recruitment and catch advice used for management as is the ultimate goal of the ESP process (Shotwell et al., 2023; Shotwell et al., 2025). This process is explored further in Champagnat et al. (2026).

Finally, our third case study features the use of causal inference to evaluate competing mechanistic hypotheses regarding the underlying drivers of fish productivity and generate information directly relevant for fishery managers and other interested parties. We first developed DAGs based on existing literature and expert knowledge, used DSEM to test DAG-data consistency (d-separation test) and quantify linkages and pathways that may drive fish productivity, and compared support for different causal hypotheses. EBFM calls for integrating the physical, biological, socioeconomic, and population components of an ecosystem into management (Link, 2002; Pikitch et al., 2004). DSEM provides a flexible approach to explicitly model how these components, and, crucially, their interactions, drive dynamics of populations over time (Thorson et al., 2024). As illustrated above, this can include the incorporation of lagged effects, allowing formal evaluation of hypotheses that link spawning stock condition and biomass to subsequent recruitment (Perretti et al., 2017). Through this exercise, we identified that the drivers of abundance of young fish were driven more by prey availability compared to demography, but the most supported causal driver did vary with population and within regions (and some populations had evidence for more than one causal driver). Without DSEM, testing the competing hypotheses of prey availability and demography - which included indirect, direct, and lagged effects - would have required a sequence of separate linear models that would then obscure the relative influence of different mechanisms under study (Arif & MacNeil, 2023; Byrnes & Dee, 2025; Grace & Irvine, 2020). Identifying mechanisms governing the abundance and productivity of early life stages provides an early indicator of future stock trajectories, improves forecasts of population dynamics, and informs management of target stocks and broader ecosystem components (Houde, 2008). More broadly, this case study illustrates how DSEM can be used to not only identify causal drivers of ecosystem dynamics but how to distinguish among competing mechanistic pathways associated with different hypotheses.

This work reveals several best practices for using DSEM to support EBFM and, more broadly, to understand the dynamics of complex systems. First, in all applications, it is highly recommended that the inferential objective be identified as this will guide the DSEM workflow from the development of graphical models to model specification and selection of validation steps. In the first case study, our goal was to predict a high-level summary of system trends, e.g., indicators of benthic and pelagic biomass using an analysis that avoided shrinking species towards a shared trend. Note that we did not conduct model selection because the goal was not to provide a parsimonious description of system dynamics. Similarly, we did not use a d-separation test to validate the model because the model was not intended to provide a causal representation of system responses under a hypothetical intervention (the goal of causal analysis). Finally, we did not interpret the effects of individual paths in the DAG. The goal of the second case study was to maximize predictive power rather than making strong causal statements about the ecosystem or testing alternative hypothesized causal structures. This involved developing a DAG from expert knowledge and the literature, using DSEM to fit the structural model and a few for comparison (iid, ar1, regression models), and comparing support for different models using AIC and the proportion of unexplained variance. The last case study illustrated how DSEM can be used to compare support for hypothesized causal structures of ecosystem dynamics and make causal statements about relationships. Doing so involved setting up multiple DAGs that represent different hypotheses regarding the underlying structure, ensuring the DAGs are consistent with the data-generating process (d-separation tests), using DSEM to estimate parameter estimates, and comparing statistical support for each model using AIC (Thorson et al., 2024). Another important best practice is collaboratively developing a graphical model (or a few depending on inferential objective) based on knowledge from multiple sources including the literature, theory, and expert opinion, then transparently reporting these assumptions (Arif & MacNeil, 2023; Grace & Irvine, 2020). In contrast, causal discovery suggests that one can purely use data to identify the most appropriate graphical model (Correia et al., 2026; Nogueira et al., 2022). We view these approaches as complementary, with causal discovery serving as one line of evidence within a broader process of causal understanding (Grace, 2026; Grace et al., 2025).

Although developing a scientifically grounded DAG is an important first step, practical guidance remains limited on managing DAG complexity so that paths can be statistically estimated and, when causal inference is the objective, assumptions can be evaluated. Many studies recommend beginning with a DAG that includes all plausible causal pathways and confounders relevant to the question at hand (Arif & MacNeil, 2023; Correia et al., 2026; Tennant et al., 2021). To reduce complexity, some studies suggest simplifying through careful consideration of prior knowledge, for particular analyses (e.g., hypothesis testing), or based on statistical significance, e.g., through causal discovery or d-separation tests (Correia et al., 2026; Pearl, 2009; Shipley, 2000). We suggest that developing and refining DAGs depends on the inferential objective and critical evaluation of the paths in a given DAG. As illustrated by the second and third case studies, a DAG may contain nonsignificant linkages that remain important for representing the biological and ecological mechanisms underlying the system; removing paths or variables solely because they are nonsignificant can bias estimates of both direct and indirect effects (Arif & MacNeil, 2023; Byrnes & Dee, 2025; Grace et al., 2010). However, modifying a DAG or revisiting the underlying data may be appropriate when effects are large and statistically significant but opposite in direction to the hypothesized relationship. In the arrowtooth flounder case study, this occurred for the relationships between prey availability and recruitment and between predation mortality and recruitment, suggesting the possible omission of variables or pathways, limitations in the available data, or both (Shipley, 2000; Tennant et al., 2021). For example, the euphausiid abundance time series used to represent prey availability contained only seven observations over the 48-year model period (i.e., DSEM predicts values at each time step for as many time steps in the longest time series) and was collected within the relatively small Kodiak core survey area (Ressler, 2019), while predation mortality was represented by model-derived estimates of arrowtooth flounder biomass consumed rather than a direct measure of predation pressure. When DSEM yields an unexpected result, we therefore recommend fitting the model with and without influential time series, revisiting the DAG structure, and examining the data, model, and individual linkages in greater detail—a process we refer to as “scientific validation.” Such iteration is a recommended component of structural causal modeling (Grace & Irvine, 2020) and will also be important in DSEM applications. Because methods for evaluating causal assumptions in DSEM, including d-separation tests, remain an active area of research, workflows and best practices should be revisited as the framework develops. To facilitate broader adoption in the meantime, we synthesize these recommendations in a workflow (Figure 8) for applying DSEM to support EBFM and, more broadly, causal inference.

There are several other important areas where clear guidance is lacking for how to use DSEM for data and hypotheses typical in marine ecosystems. One is which tool to use to assess statistical or predictive performance, and how to apply them given the goal of the analysis. We used AIC because it has the advantage that it can be used for non-nested DSEM models and is widely used in applied statistics as an approximation of predictive accuracy and thus is interpretable and accessible. However, AIC has been criticized for state-space models like DSEM and alternatives exist (Auger-Méthé et al., 2021 and references therein). A full review of these is outside the scope here, but we emphasize the need for future research on model selection tools for DSEM specifically but recommend AIC as a starting place in the meantime. For DSEM models it is also important to note that AIC (and other information criteria) measures the predictive accuracy of all variables within a DAG, unlike regression where AIC only represents the response variable (i.e., not covariates), and this needs to be reflected in the interpretation of results. This is because DSEM tries to explain the covariance of all variables in a DAG simultaneously. Importantly, this also implies that to use AIC, all variables must be in all DAGs, even if not linked to other variables, to ensure the data is equivalent as required by the criterion. As an example, the spawner biomass and spawner condition variables are included in the prey availability DSEM model fit but not linked to each other or to the response variable (abundance of young fish) and are not shown in the DAG in Figure 6 for clarity. If an analyst is interested in a specific variable, such as recruitment (e.g., Figure 4), then an alternative approach is to calculate the reduction in unexplained variation for that variable and use that as a metric to select a model (see Table 3). This emphasizes the variable of interest and is straightforward to interpret and communicate. Finally, if the priority is projecting or forecasting, then leave-future-out cross-validation can be used to select among models (e.g., Bürkner et al., 2020), with an example application in DSEM in (Champagnat et al., 2026). These three ways of assessing statistical performance have different aims and thus the analyst needs to choose the one most appropriate for the inferential goal of the study. Another area that lacks guidance is the challenge of short time series and/or inconsistent variables. DSEM can be sensitive to these issues and analysts must recognize that even converged models may be unreliable for use. We recommend performing sensitivity tests by leaving data out and constructing DAGs without those variables to ensure results and conclusions are robust and stable. However, it would be prudent for future work to more thoroughly examine the behavior of DSEM with limited data and develop more specific recommendations for analysts in those situations. We anticipate that as DSEM becomes more common in EBFM and other applications, that guidance and recommendations will improve and mature and make the DSEM framework more accessible and transparent for analysts. Our work is a first step in this process.

Although DSEM has great potential in many applications related to EBFM, ecological modeling, and beyond, it is not without limitations. First, DSEM does not currently support nonlinear effects, which are a common occurrence in ecosystem dynamics. Empirical dynamic modeling is an established framework for nonlinear causal analysis (Munch et al., 2023; Sugihara et al., 2012) and other frameworks similar to DSEM - ‘moderated DSEM’ - that can include nonlinear effects are currently being developed (Thorson & Kristensen, 2026). Second, assuming that time series are stationary—or that the mean, variance, and autocorrelation are constant over time—has proven to be a barrier to understanding the complex relationships that characterize ecosystem and population change (Litzow et al., 2018). DSEM assumes stationarity as one slope (effect) is estimated for each relationship (or linkage) (Thorson et al., 2024). Moderated DSEM will have the flexibility to fit a ‘random-slopes’ model, which estimates an effect for each year, and thus, can deal with nonstationarity (Thorson & Kristensen, 2026). Overall, we have demonstrated a range of applications for using DSEM to estimate, describe, and test hypotheses regarding complex ecosystem dynamics and trends over time, information that is critical to support EBFM. DSEM offers a powerful and flexible framework for integrating the correlated, direct, and indirect effects of multiple ecosystem components, while handling missing data and lagged and simultaneous effects. As methodological advances in DSEM and related frameworks and calls for using causal modeling in ecology increase (Grace, 2026; Thorson & Kristensen, 2026), this approach will play an important role in supporting EBFM.

## Supporting Information

### Supplementary Tables

**Table S1.** A description of the variables for the one outcome (recruitment) and 10 covariates used in causal modeling of arrowtooth flounder recruitment. We also report the name for each variable in all of the figures (‘Figure label’), a description of the variable (‘Description’), the length of the time series (‘Years’), the source of the time series data, and the latent structure specified in the model (the assumed statistical model for each time series). For a more detailed description of each variable, please consult text and Table 7A.1 of the arrowtooth flounder Ecosystem and Socioeconomic Profile (Shotwell et al. 2025). For bottom temperature and small copepod abundance, a latent structure of AR1 produced unrealistic dynamics (negative correlation parameters). The latent structures for these two variables were subsequently set to iid (no correlation).

| Variable | Figure label | Description | Year range | Source | Latent Structure |
| --- | --- | --- | --- | --- | --- |
| Recruitment | Recruits | Predicted abundance of age-1 fish | 1977-2024 | Stock assessment model | iid |
| Bottom temperature | Bottom temperature | Average haul-specific bottom temperature (°C) from AFSC bottom trawl survey | 1984-2023 | NOAA AFSC | iid |
| Predation | Predation mortality | Estimated arrowtooth flounder biomass consumed (mt) | 1977-2023 | Research multispecies stock assessment model (Climate- Enhanced, Age-based model with Temperature-specific Trophic Linkages and Energetics [CEATTLE]) | AR1 |
| Prey for age-1 recruits | Euphausiid abundance | Acoustic abundance of euphausiids integrated over the water column for the Kodiak core survey area | 2003-2019 | NOAA AFSC | AR1 |
| Sea surface temperature | Sea surface temperature | Averaged daily sea surface temperature from February to April over core habitat area | 1985-2024 | NOAA Coral Reef Watch Program (https://coralreefwatch.noaa.gov/index.php) | AR1 |
| Larval abundance | Larval abundance | Catch per unit effort (CPUE) of arrowtooth larvae in Spring | 1981-2023 | NOAA AFSC EcoFOCI | AR1 |
| Prey for larvae | Small copepod abundance | Mean catch per m3 in Shelikof Strait and Sea Valley | 1987-2023 | NOAA AFSC | iid |
| Retention dynamics | EGOA and WGOA Eddy Kinetic Energy | Seasonal mean (averaged Feb-Apr) anomalies from sea surface height at 0.125 degree resolution in the central Gulf of Alaska within an area from 160W to 146W and between the depth contours of 200-1000 m using the Smith and Sandwell 2-arc-minute bathymetry data | 1993-2025 | Copernicus Marine and Environment Monitoring Service | AR1 |
| Condition of adult arrowtooth flounder | Adult condition | Summer stratum-biomass weighted morphometric condition of arrowtooth flounder from AFSC bottom trawl surveys | 1993-2025 | NOAA AFSC | AR1 |
| Abundance of YOY fish | Nearshore age 1 fish abundance | Abundance of small arrowtooth flounder ($\geq 100$ mm and $< 200$ mm, likely age-1) in the Central and Western Gulf of Alaska and Eastern Aleutian Islands | 1992-2023 | ADF&G | AR1 |

### Supplementary Figures

**Figure S1.**
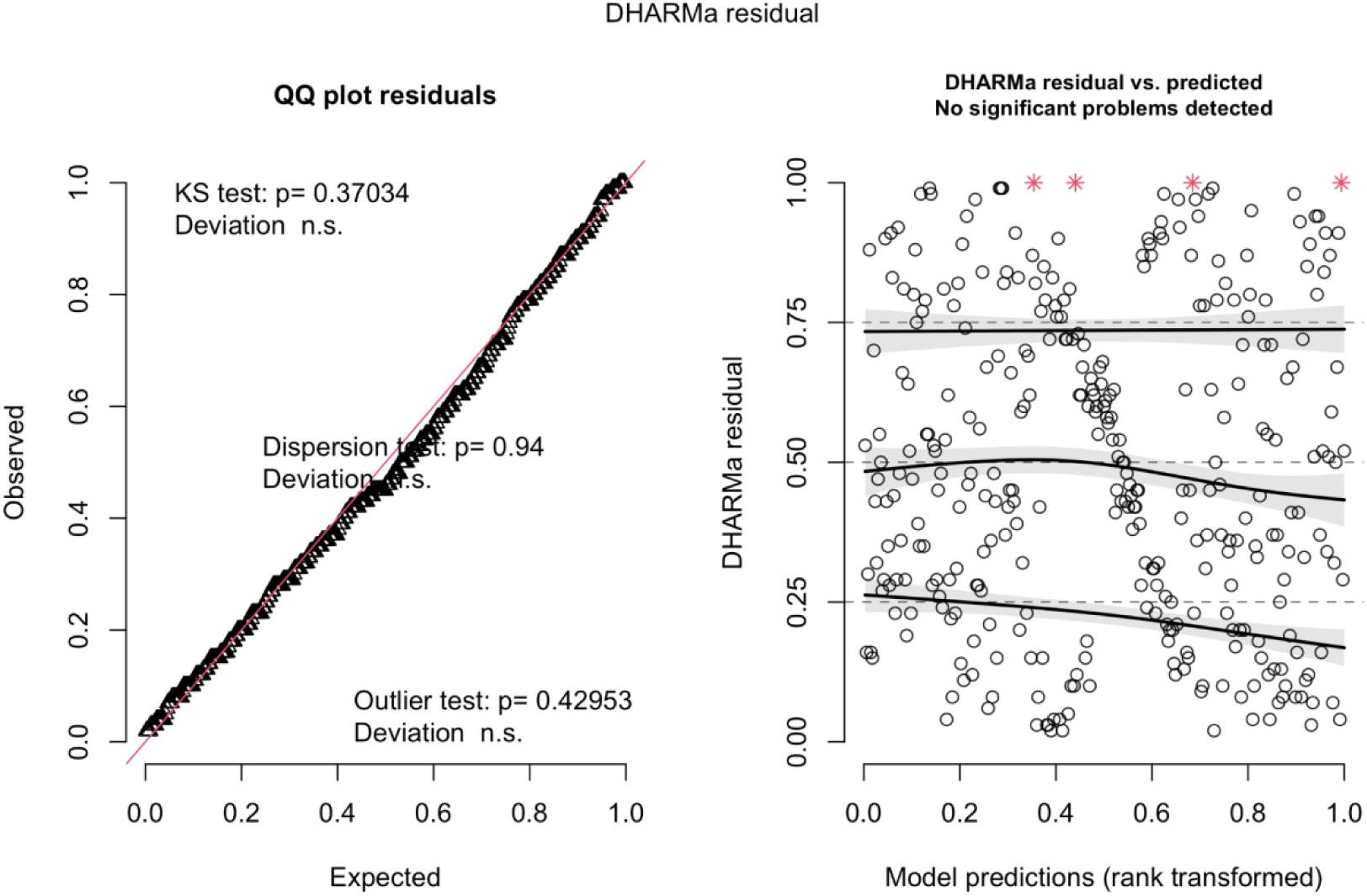
Leave-one-out residuals plotted using the DHARMa package for the arrowtooth flounder DSEM where recruitment is modeled as independent and identically distributed (iid). QQ plot (left) and residuals plotted against predictions (right). Red stars on the right plot indicate simulation outliers (data points outside range of simulated values); note probability of an outlier depends on the number of simulations (i.e., if outliers present, could be due to the number of simulations).

**Figure S2.**
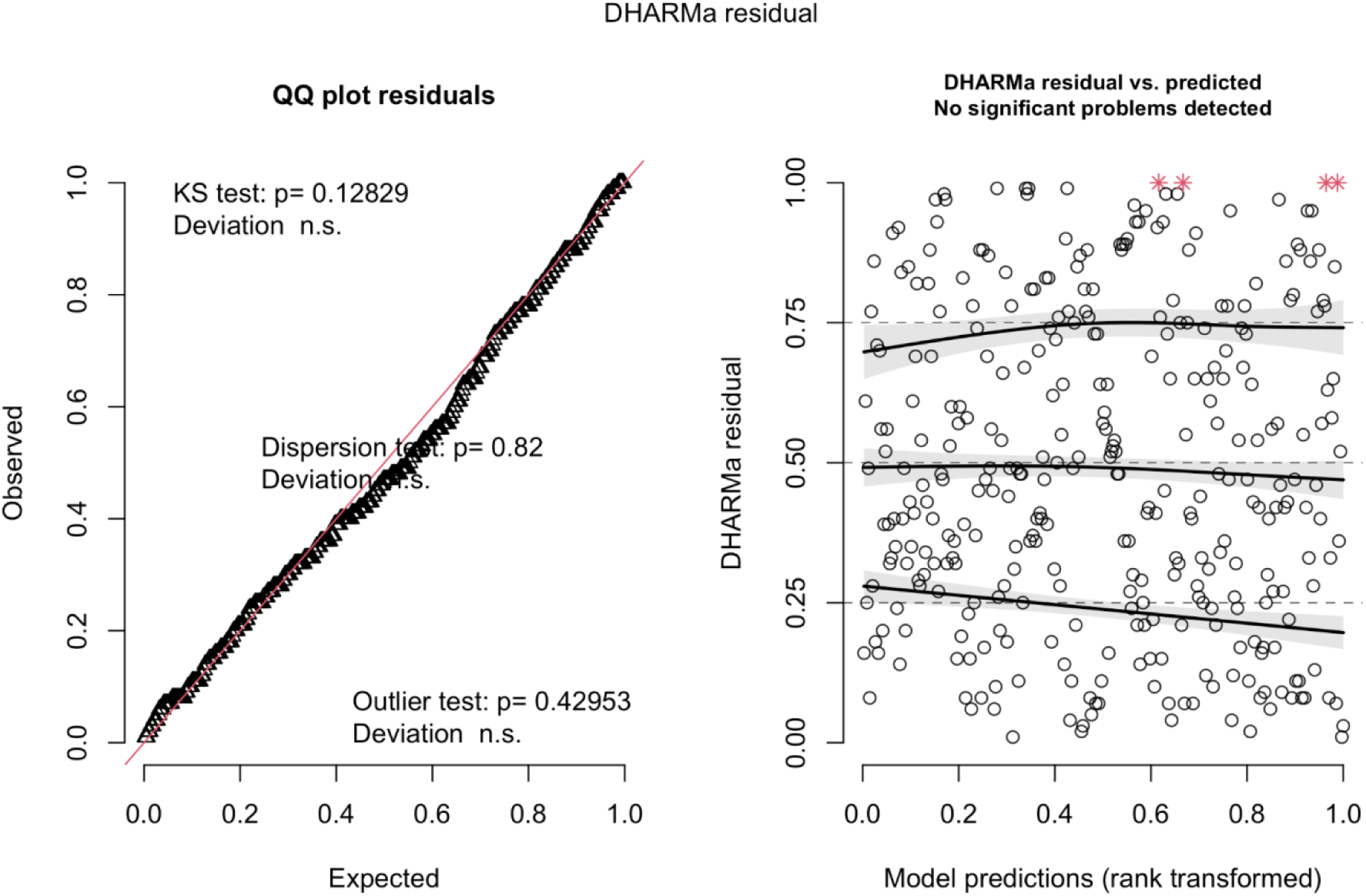
Leave-one-out residuals plotted using the DHARMa package for the arrowtooth flounder DSEM where recruitment is modeled as an autoregressive process (AR1). QQ plot (left) and residuals plotted against predictions (right). Red stars on the right plot indicate simulation outliers (data points outside range of simulated values); note probability of an outlier depends on the number of simulations (i.e., if outliers present, could be due to the number of simulations).

**Figure S3.**
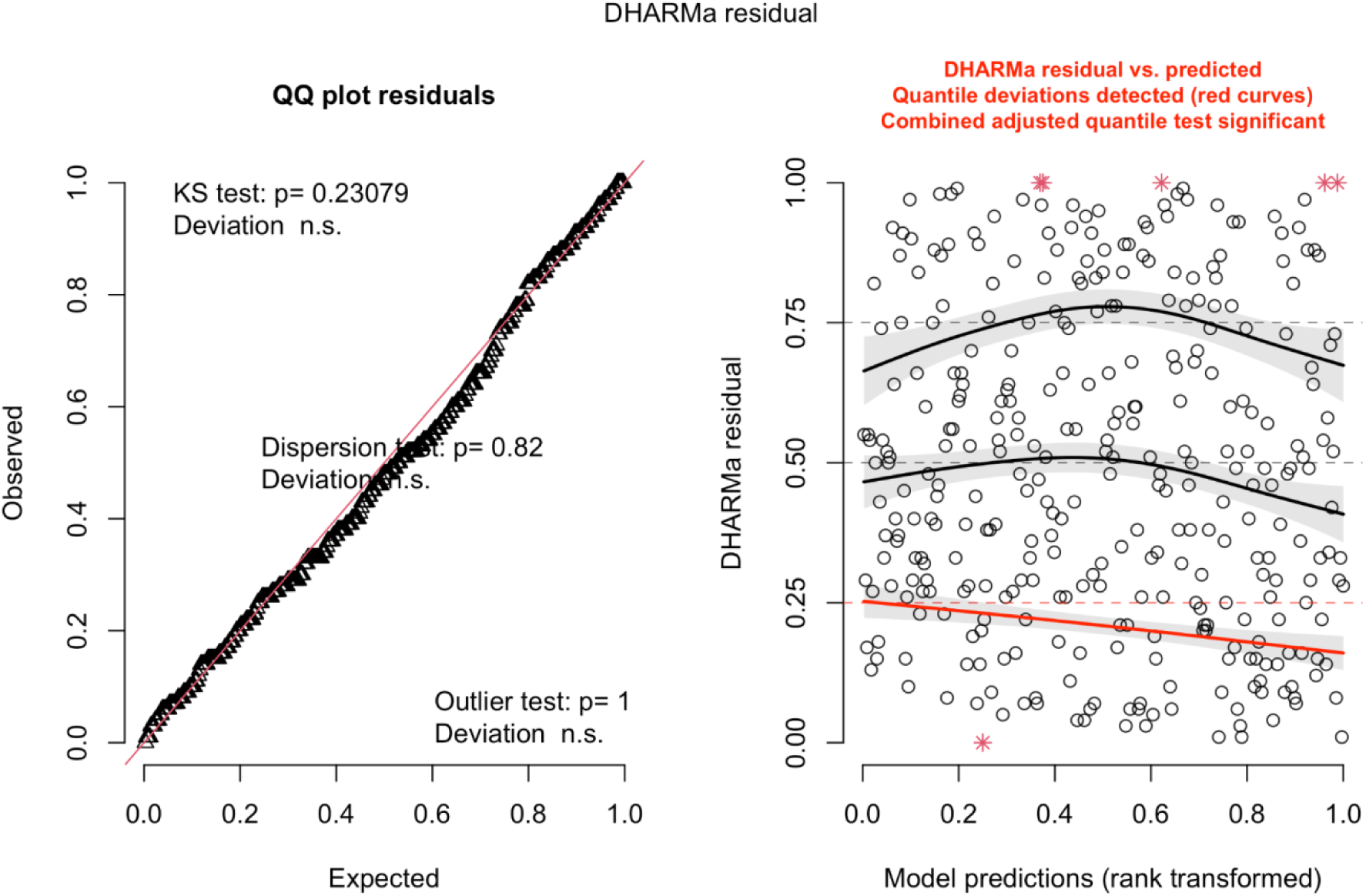
Leave-one-out residuals plotted using the DHARMa package for the regression model of the arrowtooth flounder case study where only direct links from the full causal DAG are included (see text). QQ plot (left) and residuals plotted against predictions (right). Red stars on the right plot indicate simulation outliers (data points outside range of simulated values); note probability of an outlier depends on the number of simulations (i.e., if outliers present, could be due to the number of simulations). Note while the model generally fits well, the lower quantile regression (0.25 quantile) deviates from the expected quantile, suggesting some structure in the response that is not being captured by the model. This suggests that there are missing covariates or links between covariates, nonlinear effects, or incorrect lag structures.

**Figure S4.**
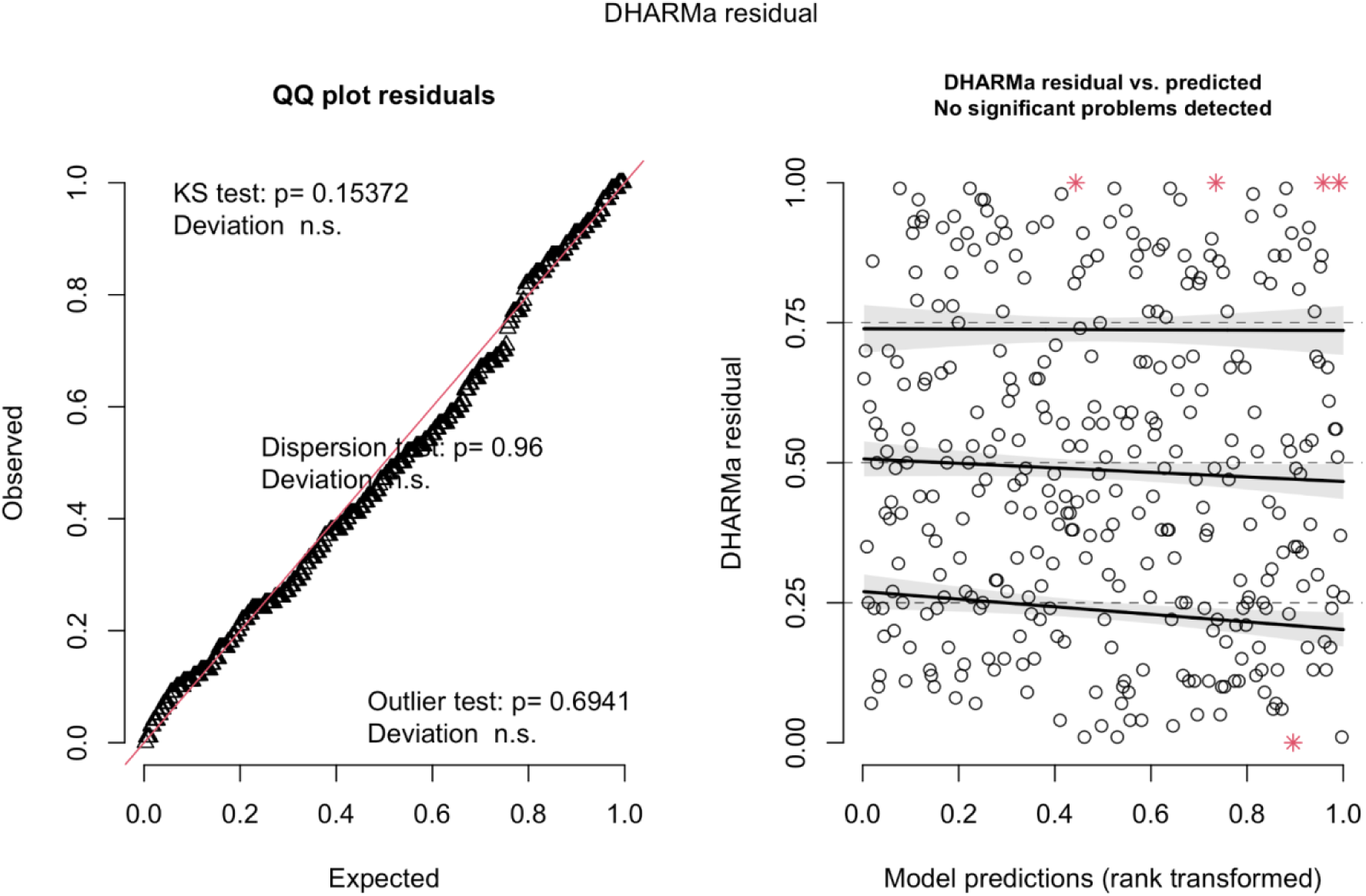
Leave-one-out residuals plotted using the DHARMa package for the causal model of the arrowtooth flounder case study (see text). QQ plot (left) and residuals plotted against predictions (right). Red stars on the right plot indicate simulation outliers (data points outside range of simulated values); note probability of an outlier depends on the number of simulations (i.e., if outliers present, could be due to the number of simulations).

**Figure S5.**
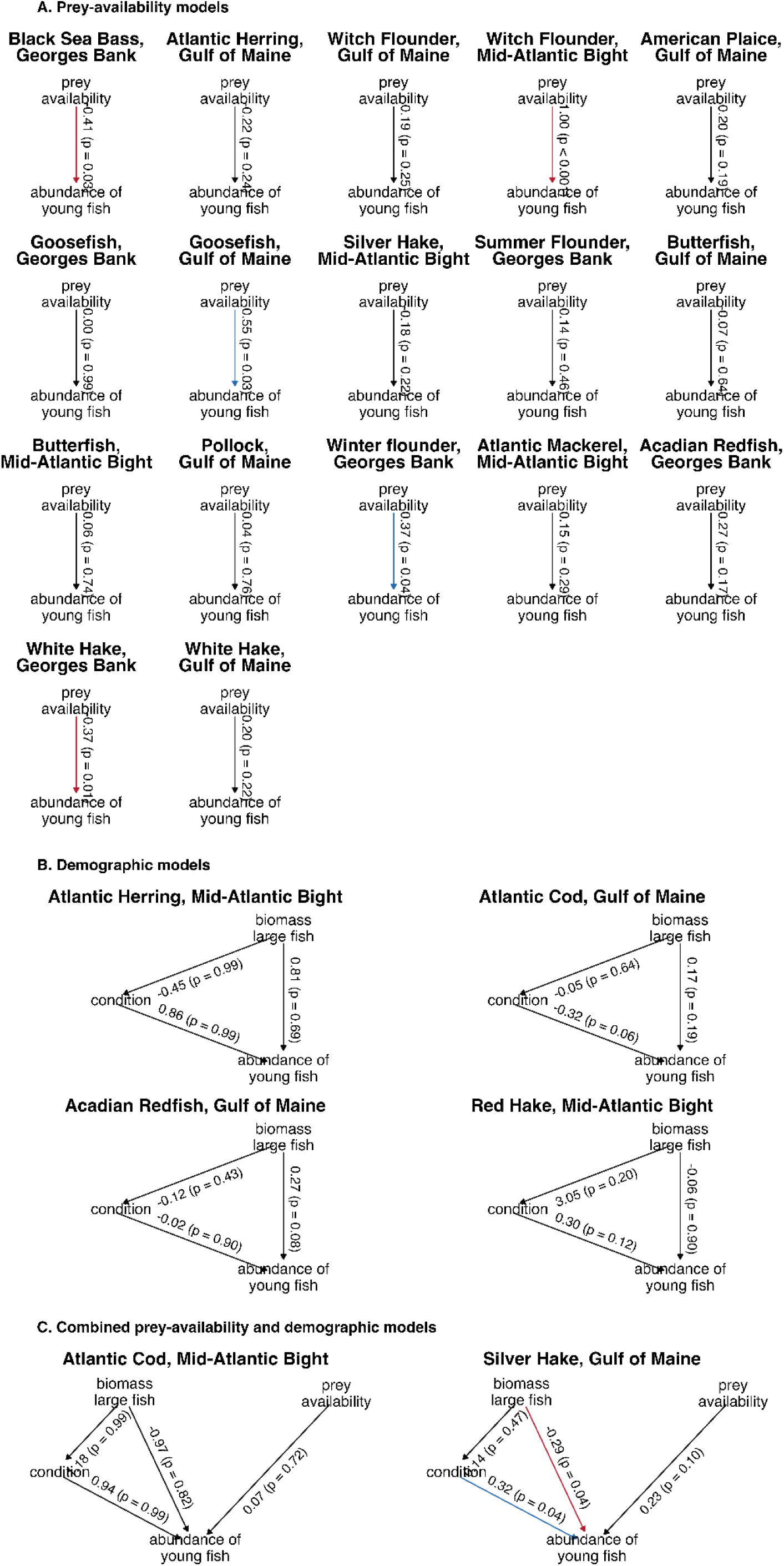
The most supported causal hypothesis that explains variation in the abundance of young fish stocks on the Northeast Shelf of the U.S. for each stock. Models are shown here for the 23 stocks that had clear support for one of the three competing hypotheses regarding abundance of young fish over time (see text, Table 4). Numbers indicate the estimated effects and *p*-values (in parentheses). The direction of the arrow indicates the hypothesized causal relationship, and the color indicates the sign (red = negative, blue = positive, black is not significant). A 1-year lag was included for the link between spawner biomass and the abundance of young fish and the link between spawner condition and abundance of young fish.

## Notes

### Competing Interest Statement

The authors have declared no competing interest.

